# Data-driven discovery of cell-type-directed network-correcting combination therapy for Alzheimer’s disease

**DOI:** 10.1101/2024.12.09.627436

**Authors:** Yaqiao Li, Carlota Pereda Serras, Jessica Blumenfeld, Min Xie, Yanxia Hao, Elise Deng, You Young Chun, Julia Holtzman, Alice An, Seo Yeon Yoon, Xinyu Tang, Antara Rao, Sarah Woldemariam, Alice Tang, Alex Zhang, Jeffrey Simms, Iris Lo, Tomiko Oskotsky, Michael J Keiser, Yadong Huang, Marina Sirota

## Abstract

Alzheimer’s disease (AD) is a multifactorial neurodegenerative disorder characterized by heterogeneous molecular changes across diverse cell types, posing significant challenges for treatment development. To address this, we introduced a cell-type-specific, multi-target drug discovery strategy grounded in human data and real-world evidence. This approach integrates single-cell transcriptomics, drug perturbation databases, and clinical records. Using this framework, letrozole and irinotecan were identified as a potential combination therapy, each targeting AD-related gene expression changes in neurons and glial cells, respectively. In an AD mouse model, this combination therapy significantly improved memory function and reduced AD-related pathologies compared to vehicle and single-drug treatments. Single-nuclei transcriptomic analysis confirmed that the therapy reversed disease-associated gene networks in a cell-type-specific manner. These results highlight the promise of cell-type-directed combination therapies in addressing multifactorial diseases like AD and lay the groundwork for precision medicine tailored to patient-specific transcriptomic and clinical profiles.

**Graphic Abstract:** 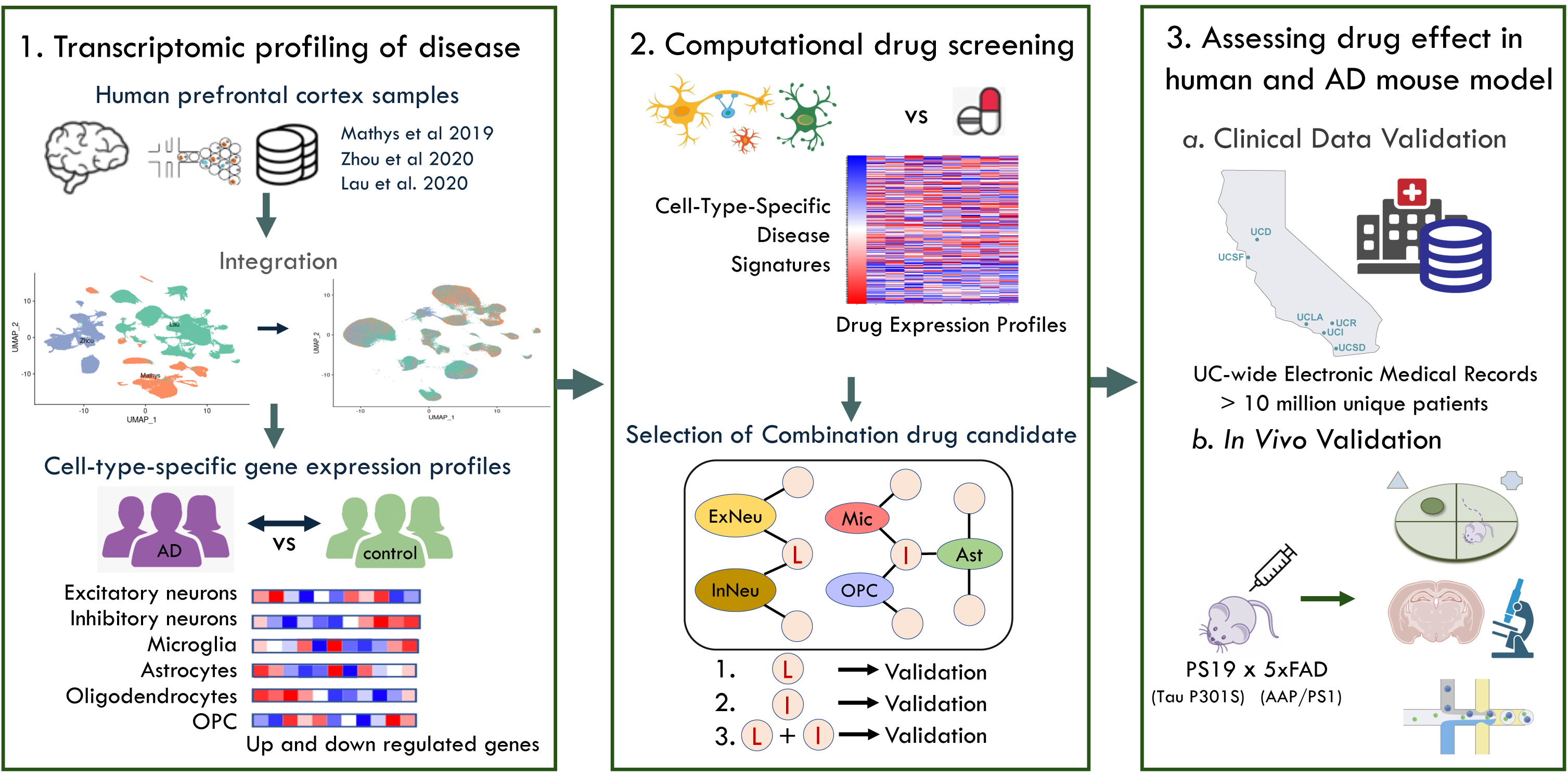

## Introduction

Alzheimer’s disease (AD) is a neurodegenerative disorder with severe impacts on individuals, families, and society^1^, and yet without a cure. AD patients suffer progressive memory loss, behavior changes, and cognitive and motor deficits, leading to diminished quality of life^2^. Over 50 million people currently live with AD or other related dementias worldwide, a number projected to triple by 2050^3,4^. With global costs exceeding $1 trillion annually, AD is one of the most costly health conditions worldwide^5^. Urgent action is needed to develop effective and accessible treatments.

Despite rigorous preclinical and clinical research efforts, AD drug development faces significant challenges, with a 98% failure rate in recent decades^6^. Current therapeutic options are mostly limited to symptom-managing treatments^7^. Furthermore, recently approved immunotherapies have only modest effects on disease progression^8,9^. The lack of effective treatments stems from the pathological heterogeneity in AD. While AD’s most prominent disease hallmark is proteopathy, characterized by extracellular amyloid-β (Aβ) plaques and intracellular tau neurofibrillary tangles (NFTs), their interplay and exact mechanisms leading to disease remain unclear^10^. Genetic heterogeneity further complicates the disease, including risk mutations in the amyloid precursor protein (*APP*) and presenilin genes, and APOE4, a risk isoform of apolipoprotein E (*APOE*)^11^. Emerging evidence highlights the critical roles of different brain cell types, with neuroinflammation and inadequate neuronal support from malfunctioning glial cells contributing to AD progression^12^. Considering the multifactorial nature of AD, traditional therapeutic approaches focusing on single disease hallmarks or bulk-tissue-level pathologies are often insufficient, leading to variable treatment outcomes.

Given the unmet need for disease-modifying treatments, drug repurposing has gained interest due to its faster development, lower costs, and improved safety^13^. In addition, technical advancements in mining large-scale databases offer new opportunities for discovering promising candidates. These developments, combined with various drug screening approaches such as *in vitro*^14^ and *in vivo* assays^15^, gene signature matching^16^, network modeling^17^, machine learning^18^, and data mining^19^, have identified numerous repurposed candidates for AD over the past decade. However, the extensive array of potential candidates complicates the establishment of priorities for clinical translation^20^.

In this study, we proposed repurposing a combination of two existing drugs, letrozole^21^ and irinotecan^22^, to reverse cell-type-specific gene expression alterations across multiple distinct cell types implicated in AD. Our drug screening strategy is entirely driven by human data, utilizing large-scale omic datasets from post-mortem brains^23–25^, a drug perturbation library generated in human cell lines^26^, and clinical records encompassing millions of individuals, thereby maximizing the chance of clinical translation. We validated our predicted drug candidates through dosing experiments in an AD mouse model^27,28^, showing that the combination therapy targeting both neurons and glial cells significantly ameliorated memory deficits and AD-related pathologies compared to vehicle treatment, and outperformed single-drug treatments targeting either neurons (letrozole) or glial cells (irinotecan) alone. The successful *in vivo* validation underscores the power of multi-cell-type network-correction therapies in effectively treating AD, highlighting the promise of our human data-driven, cell-specific drug discovery approach to develop comprehensive therapies for complex diseases. Furthermore, by targeting molecular signatures and clinical features derived from real-world individuals, this study demonstrates the potential for AI-enabled precision medicine leveraging large-scale multimodal personalized measurements.

## Results

### Computational integration of large-scale single-nucleus transcriptomic datasets from multiple studies enables the identification of cell-type-specific AD signature profiles

We organized a comprehensive single nucleus RNA-sequencing (snRNA-seq) dataset by combining published data from three independent studies^23–25^(Figure 1a), covering 37 AD patients (15 females and 22 males) and 29 controls (13 females and 22 males). Samples from individuals who did not meet both CERAD^29^ and Braak^30^ criteria for AD or control were excluded from our dataset. The Uniform Manifold Approximation and Projection (UMAP) visualization reveals clustering predominantly by the datasets of origin rather than biological variations, such as cell types or disease status (Figure 1b, Figure S1a-b). After batch harmonization with the integration algorithm, technical artifacts between datasets were effectively removed (Figure 1c), and distinct cell populations clustered by cell types (Figure 1d) while no discrete separation was observed based on disease status (Figure 1e). The integrated dataset consisted of expressions of 29,120 features in 137,065 cells.

**Figure 1.**
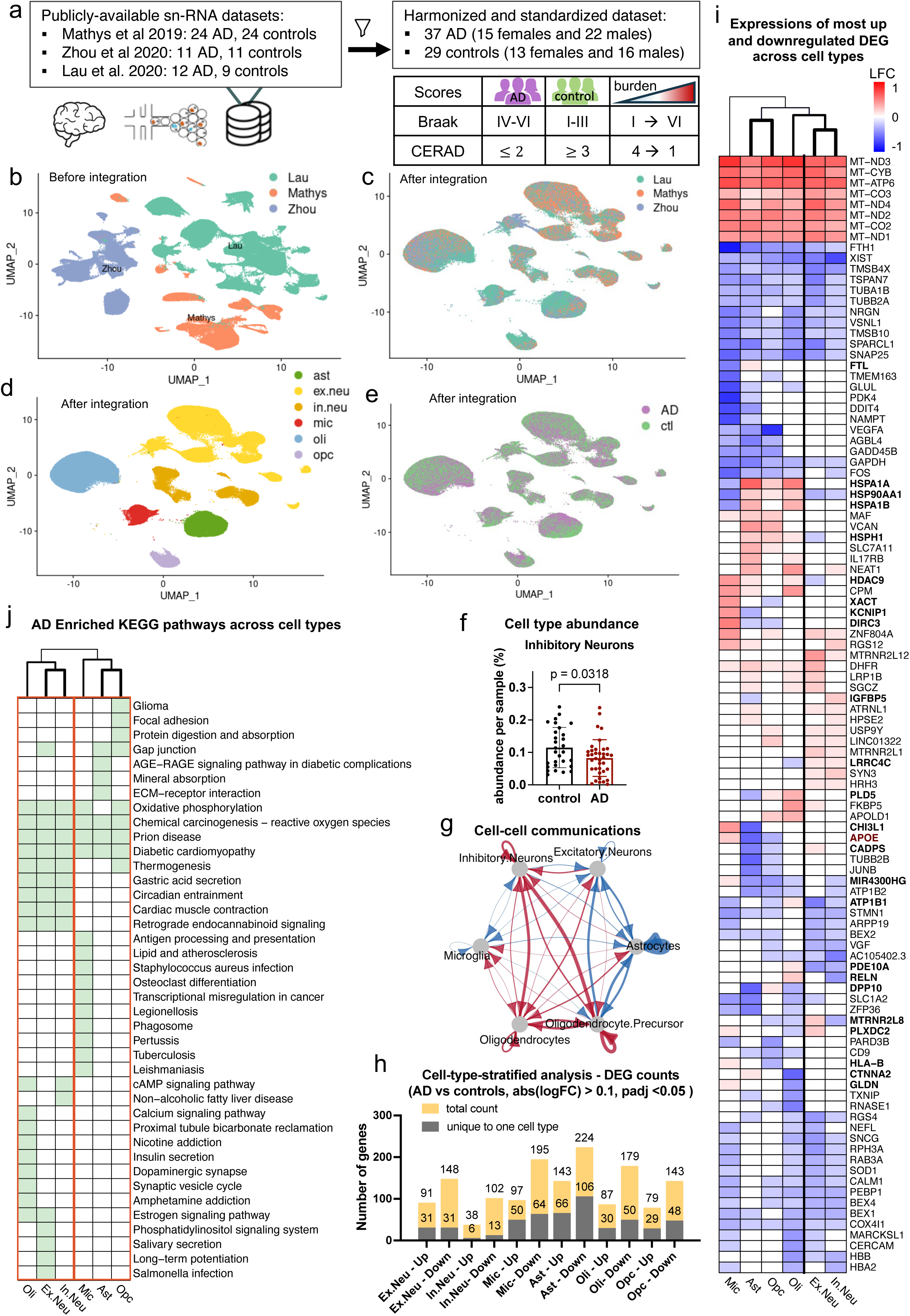
Single nucleus transcriptomic profiling reveals both shared and cell-type-specific gene expression signatures in human AD samples. a. Data sources (publicly-available) and cohort summary, including sample filtering criteria and case-control standardization strategies. b. UMAP plot of merged samples from three independent studies. Study-associated variations are apparent without integration. c-e. UMAP plots of harmonized dataset using Seurat CCA integration algorithm, with study variation eliminated, and labeled by study (c), cell type identity (d), and case-control identification (e). f. Scattered bar plot of cell type abundance percentage (values are mean ± SEM) for inhibitory neurons, comparing AD and controls (unpaired two-sided t- test). g. Projection plots for cell type to cell type communication measured by expression of receptor and ligand pairs and represented as arrows connecting two cell types. Red connections indicate increased communication and blue indicate decreased communications comparing AD vs controls. Arrows points to the directions from sender to receiver of the communication. h. Bar plot of differentially expressed gene (DEG) counts of AD versus controls in the six major brain cell types and separated into up and down regulated subgroups. Counts of DEGs unique to one cell type were superimposed on the total counts. i. DEG heatmap showing overlaps of top DEGs (highest absolute log fold changes) in each cell type, DEGs with opposite regulatory patterns in AD across cell types are bolded. j. Heatmap showing AD enriched KEGG pathways across cell types. Few functional pathways are shared by all six cell types. Oligodendrocytes cluster with excitatory and inhibitory neurons, suggesting more similarity among these cell types, while microglia cluster with astrocytes and OPCs.

To systematically characterize cell-type-specific AD pathophysiological features, we conducted comprehensive analyses focusing on six disease-relevant cell types: excitatory neurons (ex.neu), inhibitory neurons (in.neu) microglia (mic), astrocytes (ast), oligodendrocytes (oli), and oligodendrocyte precursor cells (OPC). In AD patients, the proportion of inhibitory neurons significantly decreased compared to controls (Figure 1f), as previously reported in human and mouse models of AD^31–33^. Cell-cell-communication analysis revealed heterogeneous signaling patterns among neuronal subpopulations in AD compared to controls. Signaling to inhibitory neurons increased from all cell types except microglia, while signaling to excitatory neurons decreased from all cell types except oligodendrocytes (Figure 1g).

Cell-type-specific differential gene expression analysis between AD and control groups revealed significant variations in differentially expressed genes (DEGs) across cell types, with highest number of DEGs unique to one cell type observed in astrocytes (Figure 1h). Many DEGs were unique to specific cell types, while others were shared but displayed opposite regulatory patterns in AD (Figure 1i). For example, the *APOE* gene, a major genetic risk factor for AD^34^, was significantly upregulated in microglia but downregulated in astrocytes and OPCs in AD samples. Additionally, gene set enrichment analysis showed that AD-enriched Kyoto Encyclopedia of Genes and Genomes (KEGG) pathways and gene ontology (GO) terms exhibited extensive cell-type heterogeneity, with most pathways and terms unique to individual cell types (Figure 1j, Figure S2). While pathways like chemical carcinogenesis- reactive oxygen species and prion disease were enriched in AD across all six cell types, unique pathways delineated them into two clusters. The neuronal-centric cluster included excitatory and inhibitory neurons, with oligodendrocytes sharing estrogen signaling with excitatory neurons and cAMP signaling with inhibitory neurons. The glial-centric cluster was comprised of astrocytes, microglia, and OPCs, with astrocytes and OPCs sharing gap junction pathway enrichment, and microglia and OPCs enriched for oxidative phosphorylation. These findings suggest AD pathogenesis involves heterogeneous transcriptomic-driven molecular alterations manifested in discordant behaviors of multiple cell types. Effective treatment likely needs to correct malfunctions in multiple, if not all, cell types.

### Computational drug repurposing pipeline predicts cell-type-specific therapeutic candidates

After establishing cell-type-specific AD profiles from our integrated human dataset, we screened for network-correcting drug candidates targeting AD-specific transcriptomic changes across multiple cell types. We queried each cell-type-specific AD transcriptomic profile against the Connectivity Map (CMap) drug expression database^26^ using a computational pipeline that matches gene expression profiles of diseases and existing drugs (Figure 2a). Most AD signatures overlapped with drug profile features, which were then used as inputs in the pattern-matching algorithm (Figure 2b). With a false discovery rate (FDR) < 0.05, we predicted 35 hits for excitatory neurons, 12 hits for inhibitory neurons, 8 hits for microglia, 33 hits for astrocytes, 4 hits for oligodendrocytes, and 8 hits for OPCs. Several drug hits overlapped across cell types, forming linkages in a network visualization of drug interactions with cell types (Figure 2c).

**Figure 2.**
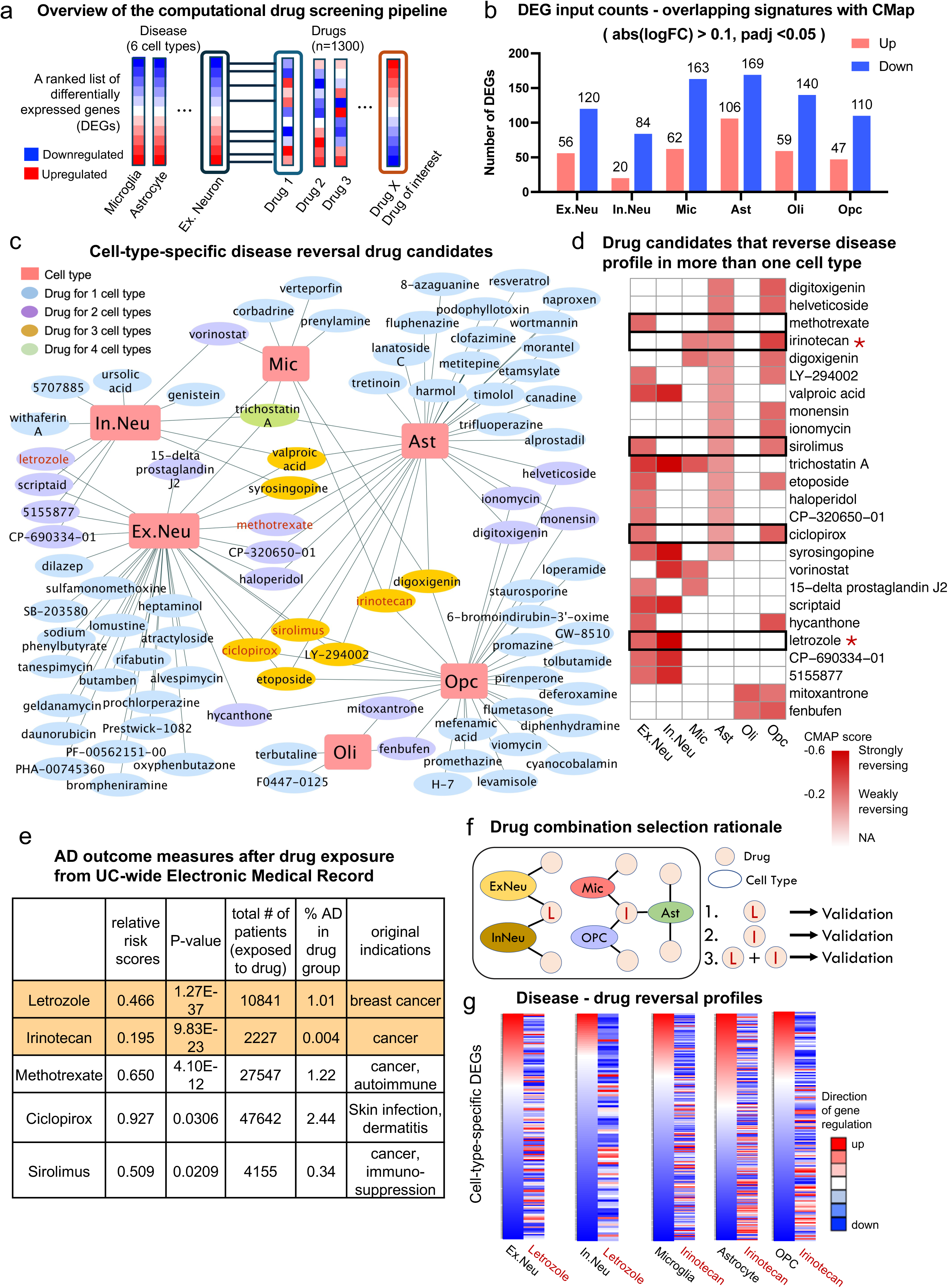
Computational drug repurposing pipeline predicts drug candidates reversing cell-type-specific transcriptomic signature profiles of AD. a, Schematic showing computational drug repurposing workflow. **b,** Input DEG counts (filtered list by mapping AD vs control DEGs to existing gene probes in the CMap database) per cell type for the computational drug repurposing pipeline. **c,** Disease-drug network depicting the connections between the six cell types and drug candidates that significantly reverse disease profiles within the respective cell types. Drug names in red denote drug candidates validated in humans using electronic medical records (EMR)**. d.** Heatmap showing drug candidates that significantly reverse AD profiles in more than one cell type. Black frames label drugs validated in humans using the EMR, and red asteroids label drugs selected for validation *in vivo* using a mouse model of AD. **e,** AD prevalence assessment in drug-exposed individuals, only drugs with significant reduced AD risks are shown. **f,** Schematic illustration demonstrating the rationale of prioritizing letrozole and irinotecan as potential combination therapy for further validation in a mouse model of AD. **g,** Heatmaps of each cell-type-specific AD transcriptomic signature profiles, rank ordered genes from the most upregulated to the most downregulated and color coded by log fold changes, in comparisons with the gene probe ranks by letrozole or irinotecan treatments, colored by corresponding fold change ranks in the CMap.

Notably, 25 repurposed drugs significantly reversed cell-type-specific AD profiles in multiple cell types (Figure 2d), indicating a multi-targeted potential with these drugs. These multi-cell-type drug candidates span various therapeutic classes, including cardiac glycosides (digitoxigenin, digoxigenin, helveticoside), chemotherapeutic agents (methotrexate, irinotecan, etoposide, mitoxantrone), aromatase inhibitors (letrozole), histone deacetylase inhibitors (trichostatinA, vorinostat, scriptaid), immunosuppressants/mTOR inhibitors (sirolimus), anti-inflammatory/anti-tumor agents (15-delta prostaglandin J2), nonsteroidal anti-inflammatory drugs (fenbufen), antifungal agents (ciclopirox), antibiotic (monensin, ionomycin), antiepileptics (valproic acid), antipsychotics (haloperidol), and several experimental compounds with undefined pharmacological classifications (LY-294002, CP-320650-01, CP-690334-01, syrosingopine, hycanthone, 5155877)^35^.

### Validation of candidate drug effects in humans using real-world evidence derived from electronic medical records

We aimed to validate the effects of the repurposed drug candidates in humans using real-world data. A distinct advantage of existing pharmaceutical agents is their potential for population-level analysis using real-world patient data to explore associations between drug use and AD outcomes. We explored the University of California (UC)-wide Electronic Medical Records (EMR) database, which includes clinical records of more than 10 million individuals across six university health centers in California. We identified 25,257 individuals diagnosed with AD within the UC-wide EMR database, out of 1.4 million individuals aged 65 or older, creating a substantial clinical dataset for analysis.

Focusing on the 25 multi-cell-type candidate drugs, we found usage records for 10 of these drugs, as only a subset of the drugs from CMap is FDA-approved or prescribed (Figure 2e, Supplementary Table 4). Of these ten drugs, five (letrozole, irinotecan, methotrexate, ciclopirox, and sirolimus) were associated with a significantly reduced risk of AD compared to matched controls, suggesting potential protective effects of them against AD. Although three drugs (etoposide, trifluoperazine, and vorinostat) showed reduced risk, statistical significance was not achieved due to insufficient patient representations. Lastly, two drugs (valproic acid and haloperidol), both used for neurological conditions, showed higher relative risk scores.

### Prioritization of letrozole and irinotecan as combination therapy for AD based on human-derived evidence

We hypothesized that combination therapy targeting both neuronal and glial transcriptomic profiles might more effectively alleviate AD pathologies. Among five drug candidates showing significant AD risk reduction in UC-wide EMR, we prioritized letrozole for its predicted reversal effects in excitatory and inhibitory neurons, and irinotecan for its effects on the glial-centric cluster, including astrocytes, microglia, and OPCs (Figure 2f). Therefore, a combination of letrozole and irinotecan potentially targets five cell types in AD.

When visualizing AD and drug profiles side by side, genes upregulated in AD neurons shifted downward, while genes downregulated shifted upward in letrozole-treated gene expression profiles (Figure 2g). Irinotecan-treated profiles showed similar shifts across glial cells (Figure 2g). Letrozole, an aromatase inhibitor primarily prescribed for breast cancer treatment and occasionally for male infertility^21,36^, had a relative risk ratio of 0.466 for AD in UC-wide EMR. It demonstrated reversal scores of -0.37 and -0.61 for excitatory and inhibitory neurons, respectively. Despite a female-skewed cohort (33 females to 1 male), the risk reduction ratios were comparable for both sexes when analyzed individually (Supplementary Table 5). Irinotecan, a DNA topoisomerase I inhibitor used for colorectal cancer treatment^22^, showed a relative risk ratio of 0.195 for AD. It demonstrated AD reversal capabilities across all three cell types of the glial-centric clusters, with reversal scores of -0.29, -0.31, and -0.45 for astrocytes, microglia, and OPCs, respectively.

### The combination treatment with letrozole and irinotecan rescues both short-term and long-term spatial memory in a mouse model of AD

To experimentally test the efficacy of the combination therapy with letrozole and irinotecan, we conducted dosing experiments in an AD mouse model expressing mutant human APP/PS1 and tau (5xFAD^28^ x PS19^27^), which recapitulates many AD-related phenotypes including amyloid plaque formation, tau tangles, and gliosis, with an early and aggressive pathology onset^37,38^ (Figure 3a). A sex-balanced double transgenic cohort was evenly divided into four groups (n=20, including both sexes) and each treated with either vehicle, letrozole, irinotecan, or the combination of both drugs every other day for three months. Spatial learning and memory performance were evaluated using the Morris Water Maze (MWM) test^39^. While no significant difference was observed in hidden platform training trials across 6 days between each treatment and vehicle group (Figure 3b), the memory test in probe trials revealed that only the combination-treatment group exhibited a statistically significant preference for the target quadrant at both 24- and 72-hours post-training (Figure 3c, d), suggesting a rescue of both short-term and long-term memory deficits by the combination treatment. Furthermore, only the combination-treatment group demonstrated significantly better location recall by making more frequent crossings to the platform location at both time points after platform removal (Figure 3e, f). Mice in the combination-treatment group also significantly outperformed the vehicle mice in latency to the first platform crossing during the 72-hour probe trial (Figure S3a, b). Average swim speeds were similar across all groups, indicating that the observed behavioral differences were not confounded by visual or motor impairments (Figure S3c, d).

**Figure 3.**
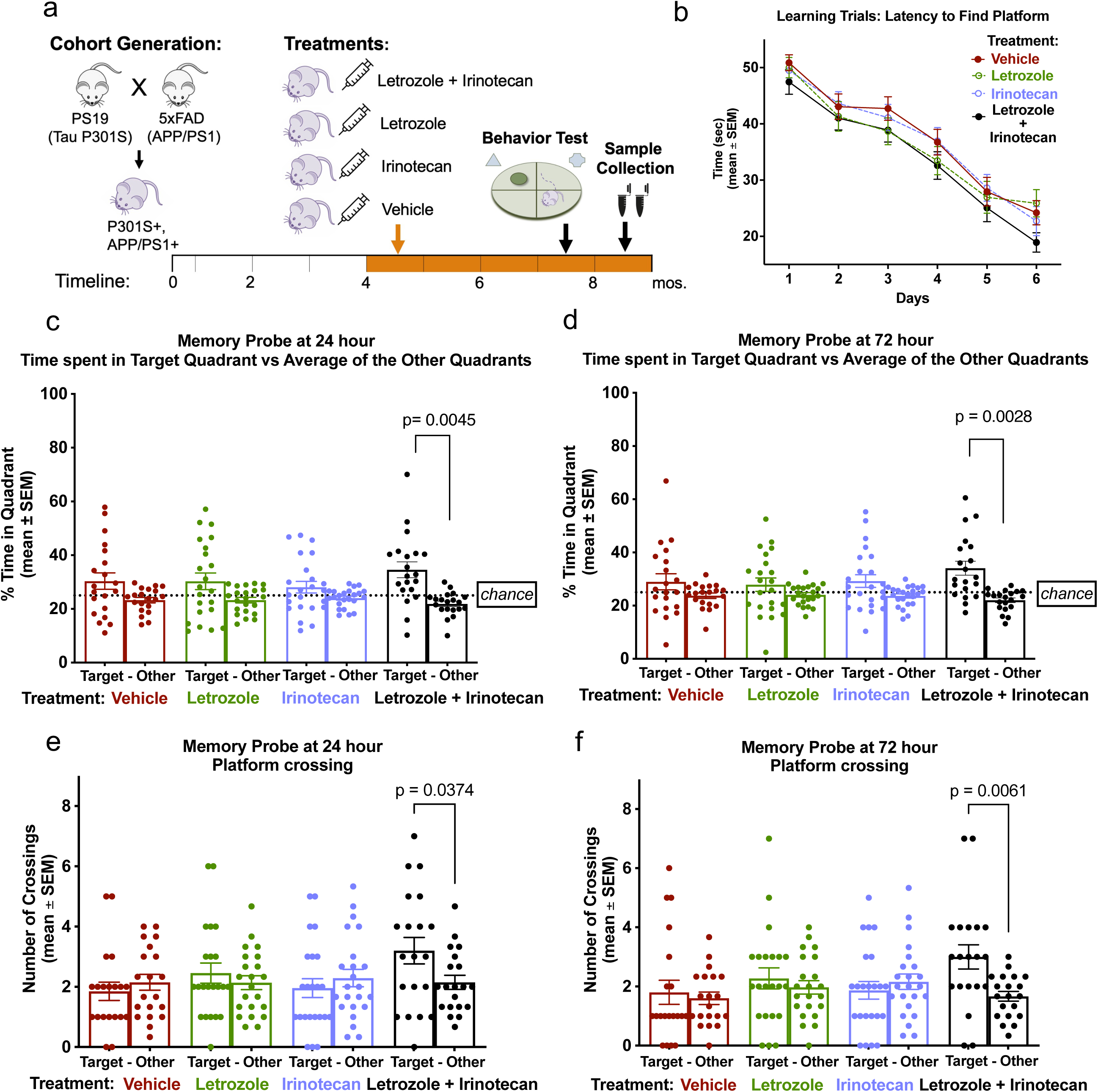
Combination treatment with letrozole and irinotecan rescues AD-like memory impairments in aged 5xFAD/PS19 mice with both Ab and tau pathologies. **a.** The schematic of treatment cohort design and experimental timeline (Created with BioRender.com). . Four treatment groups (n=20 per group, both sexes), including treatment with vehicle, letrozole (1mg/kg), irinotecan (10mg/kg) or combination of both drugs every other day via i.p. injection. **b.** Escape latency during hidden platform training days 1–6 did not differ statistically between groups. One-way repeated-measures ANOVA test was applied to compare vehicle group with treatment groups. **c,d.** Memory probes of percent time spent in target quadrant versus average of other quadrants demonstrated a significant preference of the target quadrant solely by mice with combination treatment at 24 hour (c) and 72 hour (d) after removing the hidden platform. **e,f.** Memory probes measuring number of platform-location crossings in the target quadrant versus average of the other quadrants. Significantly more crossings in the target quadrant where the platform used to be only observed in the combination treatment group at 24 hour (e) and 72 hour (d) after the platform was removed. **c-f.** Ordinary one-way ANOVA test was performed between target and other quadrants for each group, and p-value adjusted with Bonferroni multiple-comparisons testing. All graphs plot mean values ± SEM.

Additionally, sex differences were observed in the dosing experiment, with significantly improved learning performance only observed in combination-treated males compared to vehicle-treated males, and not in females (Figure S4e-f). Memory rescue by the single drug treatments was also evident in males at 24 and 72 hours of the probe trials but not in females (Figure S4g-j). Taken together, AD-related behavioral assessments, particularly in males, demonstrated that only the combination therapy with letrozole and irinotecan effectively rescued memory deficits in an AD mouse model with both amyloid and tau pathologies. Notably, the combination treatment significantly outperformed single-drug treatments, which demonstrated markedly less pronounced efficacies.

### The combination treatment with letrozole and irinotecan rescues AD pathologies in a mouse model of AD

To determine whether the single or combination drug treatments can also rescue AD-related pathologies, we morphologically assessed the cohort at 9 months of age (4 months post-treatment). According to previous literature, this AD mouse model develops extensive neurodegeneration, Aβ deposits, hyperphosphorylated tau (p-tau), gliosis, and severe loss of CA1 neurons in the hippocampus at this age^37,38^. We first evaluated neurodegeneration in the hippocampal region, as atrophy in this region is a hallmark of AD progression. Analyses of hippocampal volume showed rescue of atrophy in all treatment groups, with the most significant improvement in the combination-treatment group compared to the control group (Figure 4a, b). Thioflavin S (Thio-S) staining for β-amyloid pathology revealed significant reductions in all treatment groups compared to vehicle-treated controls, measured by the Thio-S-positive percent area and plaque counts normalized by hippocampal size (Figure 4c, d). P-tau pathology was assessed by immunofluorescent staining using the p-tau-specific AT8 antibody. While all treatment groups showed a trend of reduced p-tau pathology compared to vehicle-treated controls, only the combination-treatment group had a statistically significant reduction in the p-tau coverage area of the hippocampus (Figure 4e, f).

**Figure 4.**
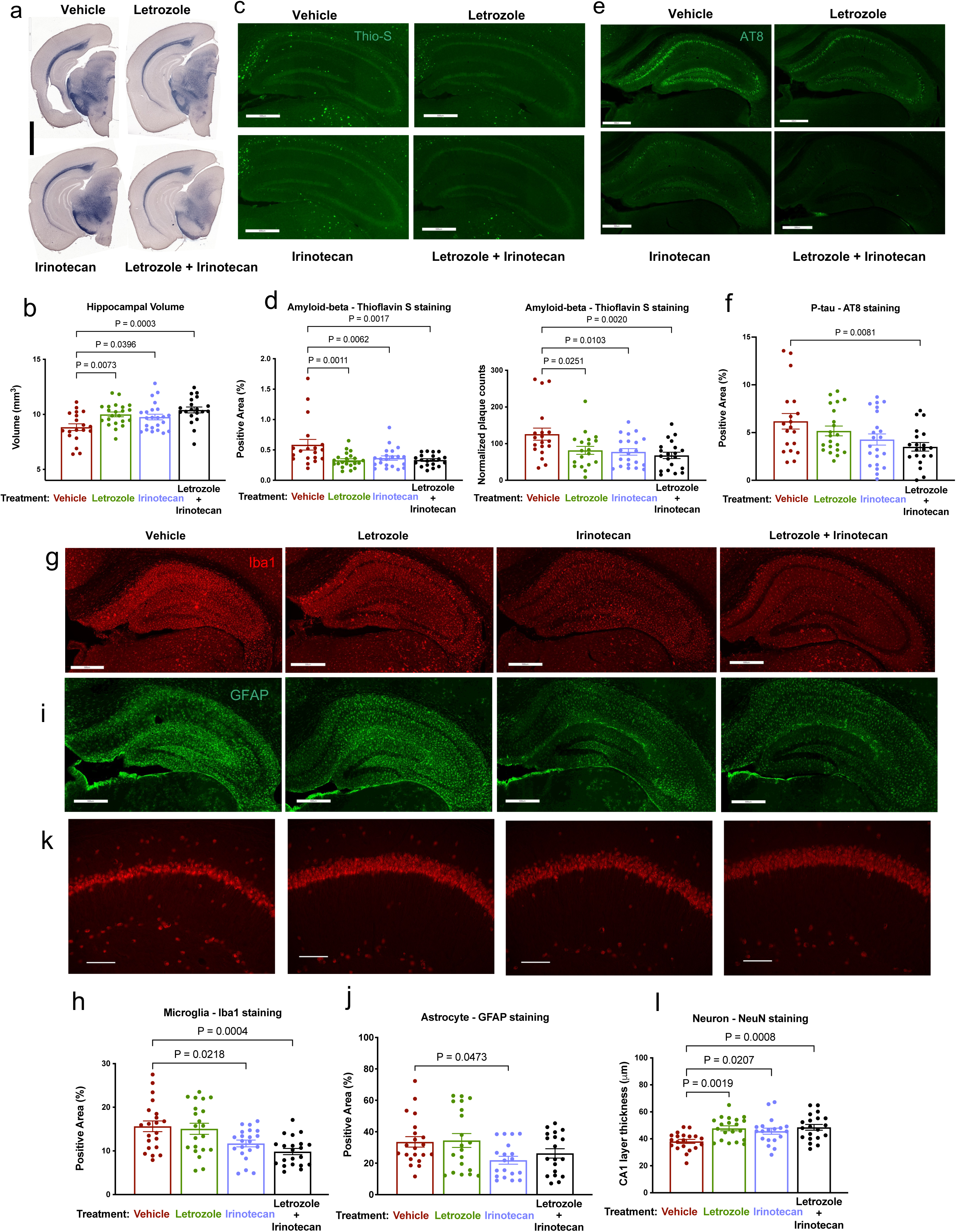
AD pathologies are significantly reduced in 9-month-old 5xFAD/PS19 mice after drug treatments, with the strongest rescue in the combination treatment group. **a.** Representative images of the ventral hippocampus from 9-month-old 5xFAD/PS19 mice with Sudan Black staining to enhance hippocampal visualization (scale bar, 2mm). **b.** Quantification of hippocampal volume in 9-month-old 5xFAD/PS19 mice across treatment groups. **c.** Representative images of hippocampus from 9-month-old 5xFAD/PS19 mice with immunostaining of phosphorylated tau (p-tau) using AT8 monoclonal antibody (scale bar, 500mm). **d.** Quantification of AT8-positive p-tau percent coverage area in 9-month-old 5xFAD/PS19 mice across treatment groups. **e.** Representative images of Thio-S staining in the hippocampus from 9-month-old 5xFAD/PS19 mice (scale bar, 500mm). **f.** Quantification of Thio-S-positive percent coverage area and plaque counts in the hippocampus of 9-month-old 5xFAD/PS19 mice across treatment groups. **g.** Representative images of microglia immunostaining with anti-Iba1 in the hippocampus of 9-month-old 5xFAD/PS19 mice (scale bar, 500mm). **h.** Quantification of the percent Iba1 coverage area in the hippocampus of 9-month-old 5xFAD/PS19 mice across treatment groups. **i.** Representative images of astrocyte immunostaining with anti-GFAP in the hippocampus of 9-month-old 5xFAD/PS19 mice (scale bar, 500mm). **j.** Quantification of percent GFAP coverage area in hippocampus of 9-month-old 5xFAD/PS19 mice . **k.** Representative images of CA1 neurons with neuronal marker NeuN immunostaining (scale bar, 100mm). **l.** Quantification of the thickness of the CA1 neuronal cell layer of 9-month-old 5xFAD/PS19 mice across treatment groups. **b, d, f, h, j, l.** Ordinary one-way ANOVA were applied between all treatment groups and vehicle control group, p-values were adjusted by Dunnett’s multiple comparison test. All graphs plot mean values ± SEM.

Given the cell-type-precision therapeutic design, we investigated the effects of each treatment on gliosis and neuronal loss through immunostaining of specific cell types. As major contributors to neuroinflammation, microgliosis and astrogliosis were evaluated by quantifying the coverage area of cell type-specific markers Iba1 and GFAP, respectively. Mice in irinotecan and combination-treatment groups showed a significant reduction in the Iba1-positive percent area in the hippocampus, indicating alleviation of microgliosis (Figure 4g, h). Astrogliosis reduction was moderate, with significant reduction observed only in the irinotecan-treated group (Figure 4i, j). Neuronal loss was assessed by measuring neuronal layer thickness via NeuN staining in the CA1 region of the hippocampus. Significant rescue of neuronal loss was evident in the CA1 region (Figure 4k, l). The most significant rescues were observed in letrozole and combination-treatment groups. Interestingly, irinotecan-treated mice also showed moderate rescue in neuronal loss, likely attributable to the alleviation of gliosis that may help prevent neuronal loss.

The combination-treatment group demonstrated improvement across all pathology markers except astrogliosis, whereas the single treatments only partially addressed these pathologies and notably failed to reduce p-tau levels. To evaluate the relationship between cognitive performance and underlying neurodegenerative changes, correlation analysis was conducted between behavioral test metrics and pathological measurements. Significant correlations were observed between memory probes, percent time spent in target quadrant, and pathology metrics, including NeuN layer thickness, hippocampal volume, p-tau level, and plaque counts (Figure S5). Notably, the combination-treatment group exhibited consistently stronger correlations compared to the vehicle-treated group, particularly between the percent time spent in the target quadrant during memory probes and the thickness of NeuN-positive layers. These findings suggest that improvements in cognitive performance in the combination-treatment group were more closely associated with reductions in pathological markers, reinforcing the potential therapeutic synergy observed.

### The combination treatment with letrozole and irinotecan promotes neuroprotective functional pathways in a cell-type-specific manner

To investigate the cell-type-specific transcriptomic changes in response to the combination treatment, snRNA-seq was performed on dissected hippocampi obtained from mice on combination treatment of letrozole and irinotecan (L+I) or vehicle treatment (n=8 per group, including both sexes). After standard processing and quality control, we obtained a filtered dataset containing 25,642 gene features expressed across 237,853 nuclei for further analysis. Through graph-based clustering and visualization using UMAP, we identified 31 distinct cell clusters, including clusters assigned to the six major cell types (Figure 5a). Additionally, we found that the L+I treatment reduced the excessive communication between inhibitory neurons and other cell types (Figure 5b, Figure S6), evident in AD (see Figure 1g).

**Figure 5.**
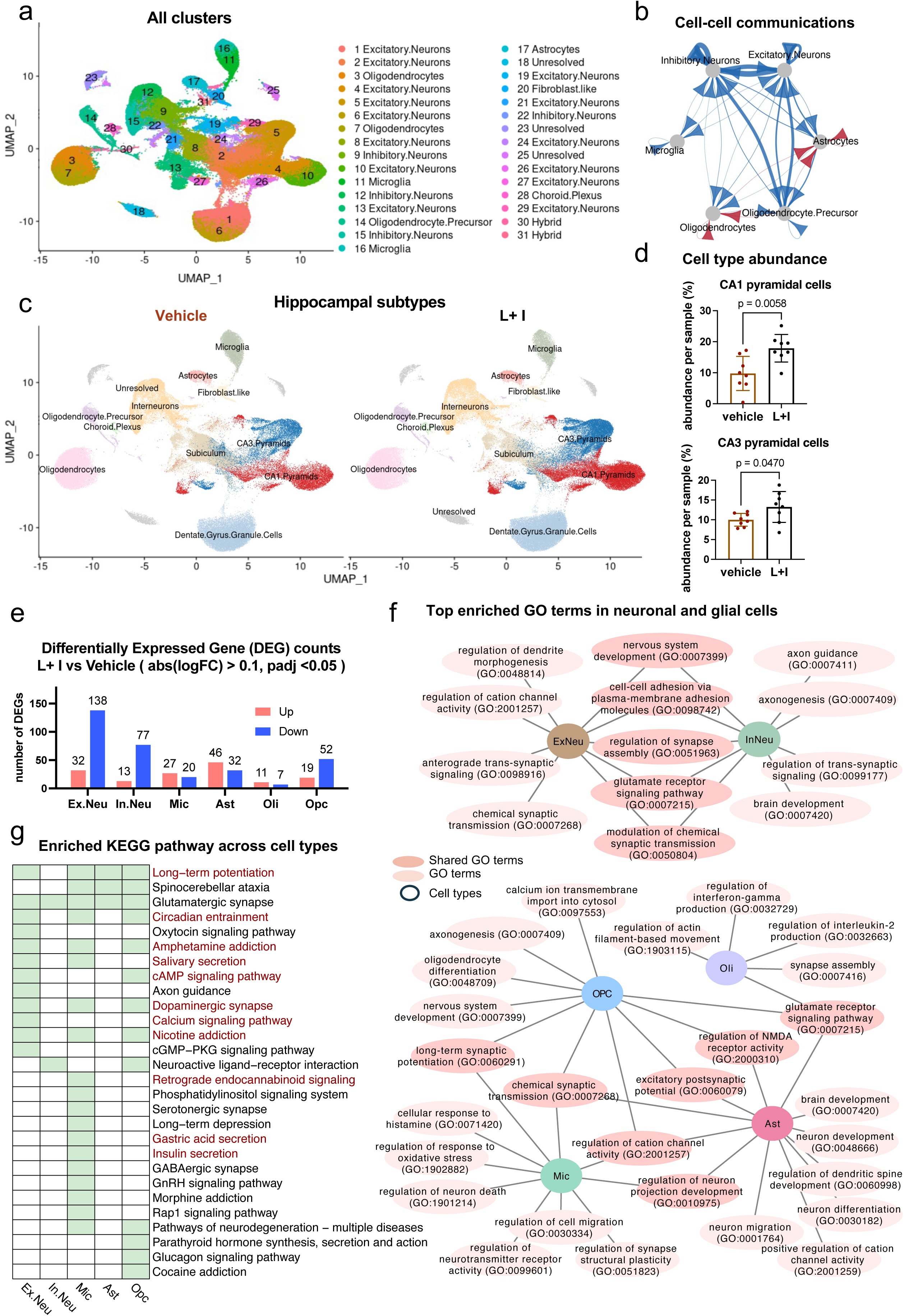
Single-nucleus RNA-sequencing (snRNA-seq) analysis in 9-month-old 5xFAD/PS19 mice across treatment groups. **a.** UMAP plot of all 31 distinct cell clusters in hippocampus of mice from combination-treatment and vehicle-treatment groups. **b.** Projection plots for cell type to cell type communication measured by expression of receptor and ligand pairs and represented as arrows connecting two cell types. Red connections indicate increased communication and blue indicate decreased communications comparing L+I-treated versus vehicle-treated groups. Arrows points to the directions from sender to receiver of the communication. **c.** UMAP plots labeled by hippocampal specific cell types and split by treatments to highlight difference in distribution density for neuronal subtypes between treatments. **d.** Scattered bar plots of cell type abundance (percentage) for CA1 and CA3 pyramidal neurons (unpaired two-sided t-test), comparing L+I-treated and vehicle-treated mice (n=8 per group, including both sexes, values are mean ± SEM). **e.** Differentially expressed gene (DEG) counts of L+I-treated versus vehicle-treated mice in six major brain cell types. Counts of DEGs per cell type do not correlate with cell counts. **f.** Top combination treatment-enriched Gene Ontology (GO) terms across six cell types. **g.** Heatmap illustration of enriched KEGG pathways across six cell types. Some treatment-enriched pathways overlap with AD signature pathways from integrated human snRNA-seq analysis (see Fig. 1) and were labeled in red.

To account for the granularity and selective vulnerabilities among neuronal subtypes, we subdivided the excitatory and inhibitory neurons into CA1 pyramidal cells, CA3 pyramidal cells, dentate granule cells (DGC), subiculum neurons, and interneurons based on expressions of hippocampal subregion marker genes. Notably, in the UMAP plot split by treatment groups, there is a discernible higher density of cells observed in the L+I-treated group compared to the vehicle-treated group within the pyramidal neuron clusters of the CA1 and CA3 regions (Figure 5c). This was quantified via cell type abundance analysis, revealing that the proportions of CA1 and CA3 pyramidal neurons in L+I-treated mice were significantly higher than in vehicle-treated mice (Figure 5d), consistent with our findings from pathological analysis.

Differential gene expression analysis comparing L+I and vehicle treatments revealed varying counts and compositions of DEGs (abs(logFC) > 0.1, padj < 0.05) across cell types, with the fewest DEGs observed in oligodendrocytes, suggesting they were least affected (Figure 5e, Supplementary Table 6). Gene set enrichment analysis of cell-type-specific DEGs revealed enrichments in disease-relevant functional pathways (Figure 5f). Notably, the treatment enriched pathways related to nervous system development and synaptic activities in neurons. For example, in excitatory neurons, the dendrite morphogenesis pathway was enriched, while the axonogenesis pathway was enriched in inhibitory neurons. In glial cells, oligodendrocyte differentiation pathway was enriched in OPCs, and actin filament-based movement and synapse assembly pathways in oligodendrocytes. Essential microglial functions like pathways of response to oxidative stress and histamine, regulation of neuron death and synaptic plasticity, and neuron projection development were enriched, as were neuron development and regulation of synaptic activity pathways in astrocytes. KEGG pathway analysis revealed that L+I-treatment perturbed pathways associated with AD, such as long-term potentiation, circadian entrainment, cAMP signaling, and calcium signaling (Figure 5g), suggesting that the combination treatment promotes neuroprotective functional pathways in a cell-type-specific manner.

### The combination treatment with letrozole and irinotecan effectively reverses multiple cell-type-specific transcriptomic signatures of AD

Lastly, we explored combination treatment-reversed transcriptomic signatures of AD to elucidate potential molecular and cellular mechanisms underlying the observed benefits of the L+I treatment. We mapped human AD signatures to homologous mouse genes. With an absolute log fold change cutoff of 0.01, the L+I treatment reversed the expression patterns of AD signatures in multiple cell types: 56 (27%) in excitatory neurons, 36 (29%) in inhibitory neurons, 90 (27%) in astrocytes, 52 (20%) in microglia, 37 (15%) in oligodendrocytes, and 64 (34%) in OPCs (Figure 6a, Figure S7). With a p-adj-value < 0.05 cutoff, 49 and 27 AD signatures were significantly reversed by L+I treatment in excitatory and inhibitory neurons, respectively, with 13 shared between them (Figure 6a). In glial cells, fewer reversed gene expressions reached statistical significance: 13 in astrocytes, 6 in microglia, 11 in oligodendrocytes, and 19 in OPCs. However, they unveiled intriguing patterns. For instance, *APOE,* identified as a DEG in the human integrated analysis, was upregulated in microglia and downregulated in astrocytes and OPCs (see Figure 1i). The expression patterns of *Apoe* in the tested AD mouse model were reversed by L+I treatment across all three cell types (Figure 6a), although statistical significance was only achieved in astrocytes and OPCs, possibly due to the presence of a small but diverse microglia population in the sequenced cohort.

**Figure 6.**
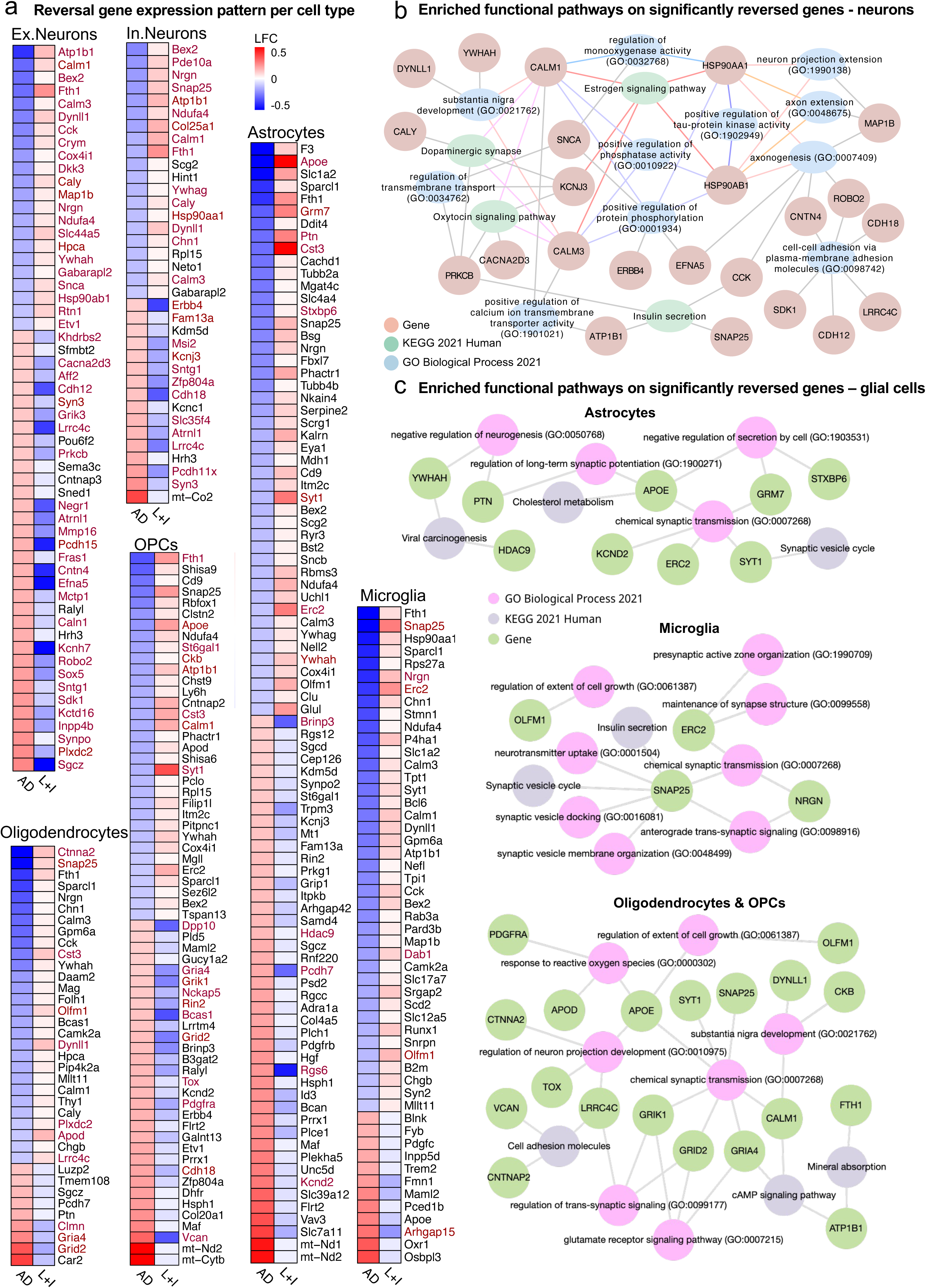
Combination-treatment with letrozole and irinotecan reverses cell-type-specific transcriptomic signatures of AD. **a.** Comparison of cell-type-specific transcriptomic signature profiles of AD in humans with gene expression changes in combination treatment versus vehicle treatment groups of mice. Only treatment- reversed genes from AD profiles are included (abs (LFC) >0.01 in L+I-treated versus vehicle-treated mice), and those with FDR-adjusted p value < 0.05 are labeled in red. Heatmap colors indicate directions and magnitude of gene expression changes, with downregulations in blue and upregulations in red. **b.** Gene set enrichment analysis on treatment-reversed genes, which reached statistical significance, in excitatory and inhibitory neurons revealed reversal of multiple AD relevant functional pathways. **c.** Gene set enrichment analysis on treatment-reversed genes, which reached statistical significance, in astrocytes, microglia, or oligodendrocyte and OPCs demonstrated reversal of glial cell specific AD gene signatures and functional pathways.

We then conducted gene set enrichment analysis on significantly reversed genes to identify potential mechanistic targets. Combining reversed genes in excitatory and inhibitory neurons revealed associations with the estrogen signaling pathway among others (Figure 6b). Reversed genes within this pathway are also involved in regulation of protein kinase and phosphatase activity, as well as tau kinase activity, highlighting potential mechanisms by which letrozole may modulate neuronal dysfunction in AD. Furthermore, certain reversed genes are associated with synaptic activities and neuron projections, potentially contributing to the rescue of neurodegeneration noted in the pathological analysis. In astrocytes, reversed genes are associated with the regulation of long-term synaptic potentiation, chemical synaptic transmission, and cholesterol metabolism (Figure 6c). In microglia, reversed genes are associated with pathways regulating synapses and cell growth (Figure 6c). Additionally, we combined reversed genes from oligodendrocytes and OPCs, revealing associations with pathways related to cell growth regulation, response to reactive oxygen species, and neuron projection regulation (Figure 6c). These findings provide a transcriptomic foundation, supporting the network correction concept that L+I combination treatment rescues AD-related behavioral and pathological deficits by rectifying complex dysregulated gene networks across multiple disease-relevant cell types.

## Discussion

This study is the first to investigate the therapeutic potential of a combination therapy addressing pathology across multiple cell types implicated in AD, with each component precisely targeting transcriptomic disease signatures specific to either neuronal or glial cells. Despite that over 2,700 clinical trials primarily targeting single core pathologies were initiated in the past two decades, only two received FDA approval^6^. The limited success of single-target approaches underscores the need for strategies addressing AD’s heterogeneous pathologies across various cell types, such as synaptic loss, neuronal death, neuroinflammation, cholesterol homeostasis alteration, energy metabolism deficit, dysfunctional phagocytosis, and demyelination^12^. To address this complexity, we designed a cell-type-directed combination therapy to correct cell-type-specific dysregulated gene networks in both neuronal and glial cells. This transformative therapeutic approach employs network-correction strategy to restore essential cellular function by targeting multiple AD-related pathways simultaneously. By reversing the expressions of hundreds of disease signature genes within distinct disease-relevant cell types, this strategy provides a more comprehensive and synergistic therapeutic approach, potentially offering greater efficiency in addressing complex disease mechanisms.

While systems biology approaches and single-cell data mining have been employed for computational AD drug repurposing^40,41^, these attempts often yield extensive lists of candidates without comprehensive validation or prioritization, thereby limiting clinical translation. To address this challenge, we leveraged real-world evidence from the UC-wide EMR to examine the prevalence of AD diagnosis in patients after drug exposure. Among the predicted drug candidates with sufficient patient representation, the majority were associated with decreased relative risk for AD, indicating potential therapeutic benefits in humans. Based on their intended cell-type-specific targets, we prioritized letrozole (targeting neurons) and irinotecan (targeting glial cells) as a combination therapy from the EMR-validated drugs. Both drugs also demonstrate the ability to cross the blood-brain barrier^42,43^, ensuring delivery to the brain for therapeutic activity. Epidemiological evidence further supports this prioritization, with studies showing reduced dementia risk among breast cancer patients treated with letrozole^44^ and decreased AD risk among colorectal cancer survivors treated with irinotecan^45^. Together, these findings provide strong clinical and mechanistic support for the potential efficacy of letrozole and irinotecan as a combination therapy for AD.

Dosing validation in the 5xFAD^28^ x PS19^27^ mouse model, characterized by pronounced Aβ and tau pathologies, revealed significant rescue of memory impairment exclusively in the combination-treatment group. The combination therapy demonstrated robust alleviation across all pathology markers except astrogliosis. In contrast, single-drug treatments only partially rescued subsets of the investigated markers, highlighting the superior therapeutic effectiveness of the combination approach. Notably, only the combination treatment significantly reduced p-tau pathology, a key predictor of cognitive function and decline in AD patients according to clinical studies^46,47^, potentially accounting for the statistically significant memory improvement observed exclusively with the combination therapy. These findings underscore the importance of targeting both neuronal and glial cell pathologies to achieve a more comprehensive and effective treatment outcome.

The network-correcting effects of the combination therapy was investigated through snRNA-seq analysis of hippocampal samples from combination-treated and vehicle-treated mice. In addition to rescuing memory deficits and pathologies, the combination treatment reversed the expression of AD signature genes enriched across diverse AD-related pathways. In neurons, these included synaptic activity, axonogenesis, neuronal projections, and protein phosphorylation regulation; in astrocytes, cholesterol metabolism and long-term synaptic potentiation; in microglia, synaptic organization and cell growth regulation; and in oligodendrocytes and OPCs, regulation of neuronal projections and synaptic signaling. These findings support the hypothesis that a targeted, multifaceted therapeutic approach confers AD-specific benefits by rectifying multiple pathological processes in a cell-type-specific manner. Further investigation is required to delineate the precise mechanisms and key pathways modulated by the combination therapy.

Despite these promising findings, several limitations of this study should be noted. First, the CMap database were measured in cancer cells and may not accurately reflect brain tissue profiles; they were used to screen drug candidates and require validations in neurological models. Relevant drug perturbation databases on different brain cell types are currently unavailable. Second, validation on animal models, though necessary, may not fully represent human biology. Significant sex differences to treatment were observed in behavior tests and pathology analyses, with male mice generally responding better behaviorally to all treatments (Figure S4). Sex differences in AD are well-documented and the impact of sex hormones in AD remain inconclusive^48^. As an aromatase inhibitor, letrozole might contribute to this sex difference. However, we observed no difference between sexes in AD prevalence after drug exposures in the UC-wide EMR sex-stratified analysis. This discrepancy suggests the observed sex differences may be specific to the mouse models, as both 5xFAD and PS19 have shown strong sex difference in prior studies^49,50^. Additionally, EMR analysis presents potential caveats, as data tend to be sparse and are not collected with specific research in mind. Two of the ten drug candidates screened using EMR data, both used in neurological disorders, showed significantly higher relative risk scores for AD, suggesting detrimental effects. However, these neurological conditions are comorbidities of AD, increasing the likelihood of exposure to these drugs in AD patients. Despite efforts to propensity-match, the control populations may not accurately represent the drug-exposed populations if relevant diagnoses are missing. Therefore, rather than using relative risk scores as a measure of drug effectiveness, we relied on EMR results to prioritize drug candidates for experimental validation.

In summary, this study introduces a transformative approach to AD drug discovery by focusing on cell-type-specific interventions and predicting multi-target drug combinations based on human-derived data. Utilizing extensive omic data from post-mortem human brains and clinical records from millions of diverse patients, our methodology provides a robust framework for identifying effective therapeutic agents through real-world evidence. Preclinical validation in an AD mouse model demonstrated that the proposed drug combination significantly improved memory deficits and alleviated AD-related pathologies, whereas the single drugs alone showed limited or no efficacy, highlighting the superior therapeutic potential of the combination treatment. Our finding that the treatment reversed AD-specific gene networks supports the multi-cellular network correction concept of this strategy and facilitates further investigation elucidating the precise mechanistic targets to advance it as a novel AD therapeutic. Besides highlighting the translational potential of the proposed drug combination, this study also exemplifies the promise of a cell-type-directed, network-correcting treatment strategy. It paves the way for precision medicine leveraging AI and large-scale patient-level data. Integrating advanced computation with evolutionary scale multi-omic, clinical, and drug perturbation databases, future efforts can develop AI systems to design tailored treatments for patients based on molecular signatures and clinical profiles of specific subpopulations or individuals.

## Methods

### Human single nuclei RNA-sequencing (snRNA-seq) data curation

To ensure a diverse representation of AD patients, we curated publicly available human snRNA-seq datasets from three independent sources. Both Mathys et al. and Zhou et al. studies were obtained from the Accelerating Medicines Partnership Alzheimer’s Disease Project (AMP-AD) Knowledge Portal (https://adknowledgeportal.org) under the Religious Order Study and Memory and Aging Project (ROSMAP). The Mathys et al. dataset is accessible through https://doi.org/10.7303/syn2580853. The Zhou et al. dataset is available under the study <u>snRNAseqAD_TREM2</u> and is also accessible through https://doi.org/10.7303/syn21125841. Only individuals without TREM2 mutations were included in our integrated dataset. The third dataset by Lau et al. was obtained from Gene Expression Omnibus (GEO) under the accession number GSE157827.

### Case-control standardization across datasets

To standardize AD identification across studies, samples were re-classified into AD or control groups based on tau tangles severity with Braak clinical staging^30^ scores and Aβ burden using Consortium to Establish a Registry for Alzheimer’s Disease (CERAD) scores^29^ as a proxy. Braak staging ranges from I to VI to represent low to high levels of tau deposition. The CERAD scoring ranges from 1 to 4 to indicate high to low Aβ burden severity. Based on the available metadata from the original studies, we defined AD cases as individuals with severe tau deposition (Braak ≥ IV) and high Aβ load (CERAD ≤ 2), and non-AD controls as individuals with low tau (Braak ≤ III) and low Aβ load (CERAD ≥ 3). Individuals with scores that did not fulfill both criteria were excluded. Our final integrated dataset consisted of 37 AD cases and 29 controls.

### Human snRNA-seq dataset integration, normalization, and batch correction

Before merging the datasets, cells deemed as poor quality were removed with the following criteria: total feature count less than 500, total features less than 250, or mitochondrial gene ratio higher than 10%. Sparse features, expressed in fewer than 10 cells, were also removed.

Simply merging datasets based on principle component analysis (PCA) reveals apparent study-associated variations. To perform batch harmonization while maximizing the conservation of disease-relevant biological variance, we performed canonical correlation analysis (CCA)^51^ using the Seurat package v4.0.4 to harmonize the merged dataset. We applied the default settings to first log-normalize the merged dataset with Seurat’s *NormalizeData* function. Then, for each dataset independently, the top 2000 variable features were identified using the *FindVariableFeatures* function with “vst” as the selection method. Integration anchors were identified with the *FindIntegrationAnchors* function based on the top variable features. Finally, a harmonized dataset consisting of 137,065 cells and 29,120 features was generated using the *IntegrateData* function in Seurat.

The data matrix was linearly transformed using Seurat’s *ScaleData* function. Dimensionality reduction through Uniform Manifold Approximation and Projection (UMAP) was performed with the *RunUMAP* function, which considers the top 30 dimensions selected from the corresponding principal component analysis (PCA) obtained by running the *RunPCA* function in Seurat.

Clustering was determined based on the first 30 PCs using the *FindNeighbors* function in Seurat, which embeds cells in a K-nearest neighbor graph based on Euclidean distance in PCA space and refines the edge weights between any two cells based on the shared overlap in their local neighborhoods. Clustering was implemented using a resolution of 0.5 in the *FindClusters* function, which applies modularity optimization techniques such as the Louvain algorithm, resulting in a set of 21 distinct clusters.

### Human cell type annotation

Cell type identities were determined by applying Seurat’s AddModuleScore function to lists of known human brain marker genes (∼ 8 per cell type) collected from PanglaoDB and referenced by Jiang et al. Cell type assignment included astrocytes, microglia, oligodendrocytes, oligodendrocyte precursor cells, endothelial cells, pericytes, and excitatory and inhibitory neurons. Each cell was assigned the corresponding cell type identity that generated the highest scores among scores for all cell types. If the highest and second highest scores of a cell were within 20% of the highest score, then the cells were deemed hybrids and excluded from further analysis. We assessed the validity of the assigned cell type identities by examining the homogeneity, distribution, and separation of cell types by clusterings in UMAP plots.

### Cell-cell communication analysis

Cell-cell communication (CCC) was calculated using the *CellChat* method^52^. Briefly, *CellChat* utilizes ligand, receptor, and cofactor expression from transcriptomic data to calculate a CCC probability. First, based on a *CellChat-*curated database of ligand-receptor interactions, differentially expressed signaling genes are used to calculate the ensemble average expression of signaling genes. Communication probability is modeled using the law of mass action, and statistically significant communications are identified using a permutation test. We then evaluated the interaction across cell types (excitatory neurons, inhibitory neurons, oligodendrocytes, oligodendrocyte precursor cells, and astrocytes). For the mouse vehicle-treatment comparison, and the human control-AD comparison, the *netVisual_diffInteraction* was used to calculate the net differential interaction strength between each of the two groups.

### Cell-type-specific differential expression analysis

For each cell type, we performed differential gene expression analysis comparing AD samples to controls using the *FindMarkers* function in the Seurat package. We set the test.use a parameter to MAST, which uses a two-part generalized linear model that models gene expression rate using linear regression and expression level using Gaussian distribution, as recommended by Mou et al^53^. in a study comparing 9 DE methods for single-cell RNA sequencing analysis. We determined differentially expressed genes (DEGs) as those with adjusted p-value < 0.05 based on Bonferroni correction using all genes in the dataset and a log2 fold change (LFC) greater than 0.1.

### Pathway analysis

Pathway enrichment analysis was conducted using g:Profiler, a web tool that performs functional over- representation analysis by mapping a given gene list to known functional information sources and detects statistically significant enriched terms. As input gene lists, we queried each cell-type-specific DEG list independently with an adjusted p-value cutoff of 0.05 to obtain significant pathway enrichment in both directions. Multiple testing correction was performed using the g:SCS algorithm, which is specifically correct for p-values obtained from GO and pathway enrichment analysis. This method is designed to correct for multiple tests that may be potentially dependent on each other due to term associations. In addition to considering Gene Ontology cellular components, biological processes, and molecular functions as supplementary data, we focused our analysis on the Kyoto Encyclopedia of Genes and Genomes (KEGG) functional pathways.

### Computational drug repurposing analysis

The computational drug repurposing algorithm, which was developed by Sirota et al. and Dudley et al., and taken from Chen et al., was applied to each disease gene signature profile using the publicly available Connectivity Map (CMap) database, consisting of treatment profiles of more than 1300 FDA-approved drugs or previously investigated compounds. The drug repurposing algorithm we employed takes two inputs: 1) an ordered list of up and down-regulated genes from individuals with disease as compared to controls and 2) the data from CMap, consisting of rank FC of each gene after drug treatment relative to vehicle controls on the same plate. The pipeline assesses the disease-drug relationship using CMap scores derived from a Kolmogorov-Smirnov test (K-S test), comparing gene expression ranks in disease and by a drug. The pipeline was adapted for single-cell analysis, where each cell-type-specific DEG signature set was overlaid with 6100 drug profiles on the 1300 drugs provided by the CMap. A drug with a strong negative CMap score indicates an opposing mechanistic relationship with the disease, suggesting therapeutic potential by reversing the regulation direction of disease signatures. The absolute values of CMap scores reflect the degrees to which the drug “flips” the signature of the disease. To address variations in input counts across cell types, significantly reversed drug profiles were identified for each cell type separately using a permutation-based approach. The false discovery rate (FDR; Benjamini-Hochberg) was calculated to adjust P-values. P-values for individual drug hits were determined by comparing reversal scores to a distribution of random scores that were generated by the permutation strategy. Negative reversal scores were deemed significant if they met the criterion of FDR < 0.05. For drugs tested multiple times (e.g., in different cell lines), the profile with the most substantial reversal (lowest negative score) was used.

### Validation in real-world human electronic medical records (EMR)

While our drug screening approach generated a total of 86 drug candidates targeting six major brain cell types, we prioritized 25 drugs that significantly reverse more than one brain cell type. This allows maximizing coverage of multiple disease-relevant cell types by combining just two drugs. The beneficial effects of the top drug hits were examined by analyzing AD prevalence (% AD diagnosis) in drug-exposed individuals and compared to propensity-matched controls, via UC-wide EMR. AD diagnosis was defined with ICD10 codes G30.0, G30.1, G30.8, and G30.9. At the time of surveying, the UC-wide EMR aggregated clinical data covering 1,441,778 individuals across six University of California (UC) campuses, encompassing more than 10 million patient records including diagnosis and medication prescriptions. For each screened drug candidate, patients prescribed or taken the drug will be identified using the medication order table and string-matching for drug names. Only individuals above the age of 65 were considered.

The matched controls were identified using a propensity score matching approach based on age, age at death (if applicable), race, ethnicity, sex, original indications (for example, breast cancer was used for Letrozole and colorectal cancer was used for Irinotecan), AD comorbidity (such as hypertension and edema), and UC center locations (including UC San Francisco, UC Davis, UC Los Angeles, UC Irvine, UC San Diego, and UC Riverside). We measured the relative risk score for AD diagnosis (not including patients with only a diagnosis of Mild Cognitive Impairment) in the drug-exposed group and compared it to the matched control groups by calculating the ratios of AD to total patients in each group. Bootstrapped χ-squared tests across 10 permutations of the iterative matching of the control group were applied to establish significance. Drugs with a significantly reduced risk of AD were included in Fig 2e. Other drugs found in the UC-wide EMR but did not yield significance or reduced risk were reported in Supplementary Table 5. Different original indications were used for matching each drug, and the ICD10 codes were listed in Supplementary Table 5.

### Drug selection rationale for validation in AD mouse model

Our integrated human single-cell transcriptomic analysis revealed two distinct pathological clusters among the six cell types analyzed. Additionally, we observed that AD inhibitory neurons exhibited the most pronounced disparity between AD cases and controls, underscoring its significance in AD pathology. These discoveries informed our prioritization of drug candidates targeting these specific clusters and accommodating inhibitory neurons for treating AD. Among a selection of five promising drug candidates that demonstrated a significant reduction in AD risk in humans, we prioritized letrozole and irinotecan, since letrozole targets the neuronal pathological cluster while irinotecan targets the glial pathological cluster. We hypothesize that the combination of letrozole and irinotecan holds synergistic therapeutic effects, with each pathological cell type cluster addressed by one EMR-validated drug candidate. Consequently, we examined AD outcomes in patients receiving both drugs under their original disease indications in the UC-wide EMR. Unfortunately, fewer than 100 patients were ever prescribed both medications, rendering statistical analysis inconclusive. To explore the synergistic effects of this combination in the context of AD, we turned to direct experimentations in animal models.

### Mouse cohort generation

The 5xFAD (6SJL) Tg mice (The Jackson Laboratory, ME, USA), overexpress both mutant human amyloid beta (A4) precursor protein 695 (APP) with the Swedish (KM670/671NL), Florida (I716V), and London (V717I) Familial Alzheimer’s Disease (FAD) mutation and human PS1 harboring two FAD mutations, M146L and L286V. These mice were crossed with the widely used tauopathy model, PS19 transgenic mice expressing P301S mutation and including four microtubule-binding domains and one N-terminal insert. A large cohort of the cross, including both sexes, was genotyped, and the littermates carrying transgenes from both lines were subsequently used for the dosing experiment.

### Drug treatments

A solution of either letrozole (1 mg/kg) or irinotecan (10 mg/kg) alone, or of the two combined was prepared weekly in 0.9% sterile saline containing 10% Tween-20 to obtain the final concentrations. Treatment dosages were established based on previously tolerated doses given during cancer studies in mice. All treatments were sonicated for 30 minutes before injections to ensure proper mixing of the drugs into the solution. The treatments were given by i.p. injection every other day to the 5xFAD x PS19 double transgenic cohort, evenly split into four groups and sex-balanced, starting 12 weeks before and continuing throughout the behavioral assessment. Mice were aged 4-5 months old before the start of the treatment and the treatments lasted about 4 months until sacrificed. Body weight was measured weekly during drug treatment; injection volume was calculated based on body weight. Control mice were injected with a matched volume of vehicle, 10% Tween in 0.9% sterile saline at pH 8.5. Injections were well tolerated and had no adverse effects on health.

### Behavioral test

Mice were housed with littermate controls. Each mouse was assigned a random number, so researchers were blinded to genotype and treatment information. Male and female mice were tested in separate rooms with the same settings and test duration to avoid olfactory cues becoming a distraction with testing a mixed sex cohort on the same equipment The Morris Water Maze pool (diameter, 122 cm) contained opaque water (21 ± 13°C) with a platform 10 cm in diameter. Mice were first trained for 6 days to locate a hidden platform submerged 1.5cm below the surface of the water using distal cues surrounding the pool. These 6 days consisted of two training sessions, 2 hours apart, each consisting of two trials with a maximum latency of 60s, with a 15-minute inter-trial interval. Entry points were changed semi-randomly between trials, but distal visual cues on the walls of the behavioral testing room remained constant throughout the test. To test for memory retention, at 24 and 72 hours after the last hidden platform training, a 60-s probe trial (platform removed) was done. The entry point for the probe trial was opposite to the target. Finally, as a control for visual acuity and motor ability, mice were tested with a cued platform where the submerged platform is indicated by a black and white striped mast 15cm high extending out the surface of the water. The platform location remained constant in the hidden platform sessions but was changed for each visible platform session. Three sessions of two trials, with a maximum latency of 60s, was performed over 2 days. Performance was monitored with an EthoVision video-tracking system (Noldus Information Technology). For the probe trials, we analyzed (1) target quadrant preference- the percent time spent in the target quadrant versus average time spent in the three other quadrants, (2) platform crossing- the number of crossings over the position of the target platform versus the average number of crossings over the equivalent positions in the three other quadrants as well as (3) escape latency- a memory probe measuring how fast mice arrive at the platform location after placement in the maze. An ANOVA was used to analyze the effect of treatment, genotype, and probe timing on percent on-target crosses.

### Histopathological analyses

One hemibrain per mouse was drop-fixed in 4% paraformaldehyde prepared in 1xPBS, washed for 24 hours in 1xPBS, then cryoprotected in 30% sucrose for 48 hours at 4°C. The fixed hemispheres were cut into 30 μm thick coronal slices using a freeze-sliding microtome (Leica). These sections were then stored in a cryoprotectant solution of 30% ethylene glycol, 30% glycerol, and 40% 1xPBS at −20 °C.

Hippocampal brain sections (ten sections per mouse spaced approximately 300 μm apart, 30-μm thick) were mounted onto microscope slides from Fisher Scientific. A 0.1% Sudan Black solution was prepared by dissolving Sudan Black powder (Sigma) in 70% ethanol (KOPTEC) and mixing it with a magnetic stirrer. After centrifuging the solution at 1,100g for 10 minutes, the supernatant was filtered using a 0.2- μm filter syringe (Thermo Scientific). The brain sections were stained with the 0.1% Sudan Black solution for 10 minutes, then washed in 70% ethanol followed by Milli-Q water. The sections were then coverslipped with ProLong Gold mounting medium (Invitrogen). All sections were imaged; eight consecutive brain sections with consistent anatomic locations from AP -1.2 to -3.6 were quantified.

For Thio-S staining, several brain sections spaced 300 μm apart were mounted onto slides, following a protocol adapted from a previous study. The sections were washed with 1× PBS-T and then incubated in a solution of 0.06% Thio-S in PBS for 8 minutes. After this incubation, the sections were washed for 1 minute in 80% ethanol and then for 5 minutes in PBS-T. The sections were counterstained with DAPI for 8 minutes, washed again with PBS-T, and coverslipped. Three sections per mouse were quantified and averaged for percent Thio-S positive area and number of plaque counts (normalized by hippocampal area size per section).

For immunofluorescent staining, several brain sections spaced approximately 300 μm apart, were washed three times with 1× PBS-T (PBS + 0.1% Tween-20) (Millipore Sigma) and then incubated for 5 minutes in boiling antigen retrieval buffer (Tris buffer, pH 7.6; TEKNOVA). After this, the sections were rinsed in PBS-T before being placed in a blocking solution composed of 5% normal donkey serum (Jackson Labs) and 0.2% Triton-X (Millipore Sigma) in 1× PBS for 1 hour at room temperature. Next, the sections were washed again in PBS-T and incubated in Mouse-on-Mouse (MOM) Blocking Buffer (one drop MOM IgG in 4 ml PBS-T) (Vector Labs) for another hour at room temperature. Following the MOM block, the sections were incubated overnight at 4°C in primary antibodies diluted to their optimal concentrations. The antibodies and their dilutions included: anti-AT8 (ms, 1:300, Invitrogen), anti-GFAP (ms, 1:500, Millipore Sigma), anti-Iba1 (rbt, 1:300, Wako), anti-NeuN (GP, 1:500, Millipore Sigma). Three sections per mouse were quantified and averaged for a positive percent area.

After the primary antibody incubation, the sections were washed in PBS-T and then incubated for 1 hour at room temperature in secondary antibodies. These included donkey anti-mouse 488 (1:1,000, Abcam), donkey anti-rabbit 594 (1:1,000, Abcam), donkey anti-guinea pig 594 (1:1,000, Jackson Immuno), and donkey anti-guinea pig 647 (1:1,000, Jackson Immuno). Subsequently, the sections were washed in PBS- T and incubated in DAPI (1:30,000 dilution in PBS-T) (Thermo Fisher) for 8 minutes at room temperature. After a final wash with PBS-T, the sections were mounted onto microscope slides (Fisher Scientific), coverslipped with ProLong Gold mounting medium (Vector Laboratories) and sealed with clear nail polish.

Images for quantifications were captured using a scanning microscope (Keyence) at magnifications of ×10 or ×20, depending on the stain. To minimize batch-to-batch variation, all samples for each stain were processed simultaneously and imaged at the same fluorescent intensity. For quantifying the percent coverage area, an optimal threshold was established for each stain in ImageJ, and all samples were quantified using this threshold. To prevent bias, researchers were blinded to the sample identities. Representative images were captured using an Aperio VERSA slide scanning microscope (Leica) at ×20 magnification.

### snRNA-seq library preparation and sequencing

The other hemibrains were dissected by brain subregions, rapidly frozen on dry ice, and kept at −80 °C. Hippocampal samples were used for single-nuclei preparation. One frozen mouse hippocampus was placed into a pre-chilled 2 mL Dounce with 1 mL of cold 1X Homogenization Buffer (1X HB) (250 mM Sucrose, 25 mM KCL, 5 mM MgCl_2_, 20 mM Tricine-KOH pH7.8, 1 mM DTT, 0.5 mM Sermidine, 0.15 mM Sermine, 0.3% NP40, 0.2 units/µL RNase inhibitor, ∼0.07 tabs/ml cOmplete Protease inhibitor). Dounce with “A” loose pestle (∼10 strokes) and then with “B” tight pestle (∼10 strokes). The homogenate was filtered using a 70 µM Flowmi strainer (Bel-Art) and transferred to a pre-chilled 2 mL LoBind tube (Fischer Scientific). Nuclei were pelleted by spinning for 5 min at 4°C at 350 RCF. The supernatant was removed, and the nuclei were resuspended in 400 µL 1X HB. Next, 400 µL of 50% Iodixanol solution was added to the nuclei and then slowly layered with 600 µL of 30% Iodixanol solution under the 25% mixture, then layered with 600 µL of 40% Iodixanol solution under the 30% mixture. The nuclei were then spun for 20 min at 4°C at 3,000RCF in a pre-chilled swinging bucket centrifuge. 200 µL of the nuclei band at the 30%-40% interface was collected and transferred to a fresh tube. Then, 800 µL of 2.5% BSA in PBS plus 0.2 units/µL of RNase inhibitor was added to the nuclei and then were spun for 10 min at 500 RCF at 4C. The nuclei were resuspended with 2% BSA in PBS plus 0.2 units/µL RNase inhibitor to reach at least 500 nuclei/µL. The nuclei were then filtered with a 40 µM Flowmi cell strainer. The nuclei were counted and then ∼13,000 nuclei per sample were loaded onto 10x Genomics Next GEM chip M. The snRNA-seq libraries were prepared using the Chromium Next GEM Single Cell 3[HT kit v3.1 (10x Genomics) according to the manufacturer’s instructions. Libraries were sequenced on an Illumina NovaSeq 6000 sequencer at the UCSF CAT sequencing core.

### Sequence alignment, filtering, and counting

The demultiplexed fastq files were processed following the procedure previously described by Zalocusky et al. In summary, the fastq files were aligned to the mouse reference genome, mm10-1.2.0, which includes introns, using the *cellranger count* function (version 4.0.0) with default parameters, as detailed in the Cell Ranger documentation. Subsequently, a single UMI count file per animal/sample was generated by the *cellranger count* function. Individual UMI count files were then combined into a single count matrix using the *merge* function in the Seurat package v4.0.4. Metadata, including age, sex, and treatment information, were added to each cell.

### Pre-processing and quality control

The count matrix was further processed with Seurat by first calculating the percentage of mitochondria genes mapped per cell. The distribution of feature count, total mapped gene count, and percentage of mitochondria genes were visualized across biological samples as violin plots, and no obvious outlier was identified. We filtered the count matrix to only include cells with higher than 250 gene features, at least 500 gene counts, and mitochondria gene percentages lower than 10%. Potential misaligned or ambiguous gene features expressed in fewer than 10 cells were also removed. These quality assurance steps resulted in a final Seurat object containing 25,642 gene features expressed by 237,853 nuclei.

### Normalization, dimensional reduction, and clustering

Count normalization and dimensionality reduction were conducted following standard procedure in the Seurat package. In brief, we performed normalization and variance stabilization with an updated version of *sctransform*, v2, and principal component analysis (PCA) with *RunPCA* (npcs = 30). Dimensional reduction through Uniform Manifold Approximation and Projection (UMAP) was performed with the *RunUMAP* function and considering the top 15 dimensions selected from the corresponding PCA.

The clustering was based on the first 15 principal components (PCs) using the *FindNeighbors* function in Seurat. This function embeds cells in a K-nearest neighbor graph, considering the Euclidean distance in PCA space and refining the edge weights between any two cells based on the shared overlap in their local neighborhoods. The clustering was obtained using the *FindClusters* function, which employs modularity optimization techniques such as the Louvain algorithm, with a resolution parameter of, resulting in a set of 35 distinct clusters.

### Mouse cell type annotation, differential gene expression, and pathway enrichment analysis

Major cell types, including astrocytes, microglia, oligodendrocytes, oligodendrocyte precursor cells, and excitatory and inhibitory neurons, were classified using mouse brain cell markers in PanglaoDB, a publicly available marker gene database. Further subdivisions of hippocampus cell types, such as CA1 and CA3 pyramidal cells, were queried against hippocampal cell-type-specific marker genes as published in hipposeq (https://hipposeq.janelia.org). Cell type identities per cluster were determined by applying Seurat’s *AddModuleScore* function to sets of mouse brain marker genes. A module score for each cell type considered was calculated per cell. Each cell was assigned the corresponding cell type identity that generated the highest scores among scores for all cell types. If the highest and second highest scores of a cell were within 20% of the highest score, then the cells were deemed hybrids and excluded from further analysis. We assessed the validity of the assigned cell type identities by examining the homogeneity, distribution, and separation of cell types by clustering in UMAP plots. Each cluster is annotated with the dominated cell type identity. Minority cell types in a cluster, defined as a cell type that accounts for less than 5% of the total counts for that cluster, were considered potential hybrid cells and excluded from further analysis.

For the cell-type-specific differential gene expression analysis, we used the same procedure and tools in the human-integrated analysis. For pathway enrichment analysis, in addition to using g:Profiler, we curated the EnrichR-KG web tool, which facilitates analysis across multiple databases and provides visual representations linking significantly enriched genes with the associated pathways and GO terms. The reversal gene-pathway network was generated and downloaded using EnrichR-KG with significantly differentially reversed genes as inputs. For each cell-type-stratified analysis, the top five KEGG pathways and GO terms were displayed in a network format.

## Resource availability

### Lead contact

Corresponding

Co-corresponding

### Materials availability

No new material was generated in this study.

### Data and code availability

The single-nucleus RNA-sequencing data in combination and vehicle-treated mice was uploaded to Figshare. https://doi.org/10.6084/m9.figshare.27041767

Code for transcriptomic data analysis and integration of multiple datasets and single cell profiling used standard packages as described in the methods and will be made available on Github upon manuscript acceptance. Code for computational drug repurposing pipeline associated with the current submission is available at https://doi.org/10.1053%2Fj.gastro.2017.02.039.

The UCSF EHR and code associated with the deidentified data analysis are available to UCSF—affiliated individuals who can contact UCSF’s Clinical and Translational Science Institute (CTSI) or the UCSF’s Information Commons team for more information. UC-wide EHR is only available to UC researchers who have completed analyses in their respective UC first and have provided justification for scaling their analyses across UC health centers. (more details at https://www.ucop.edu/uchealth/departments/center-for-data-driven-insights-and-innovations-cdi2.html or by contacting.

## Acknowledgments

This study was supported by the National Institute on Aging grants R01AG060393 to MS, R01AG057683 to MS and YH, RF1AG076647, R01AG078164, and P01AG073082 to YH, and NSF 2034836 to YL. We thank Sirota and Huang lab staff for their valuable discussions about the experimental design as well as data analyses and interpretation. We also thank Eric Chow and the staff at the UCSF Center for Advanced Technology Core for advice and support with snRNA sequencing.

## Author Contributions

YL, YH, and MS designed the study. YL performed majority of the studies and data analyses. C.P.S. performed cell-cell-communication analysis. J.B. aided immunohistochemical studies and imaging. M.X. facilitated treatment dosing experiments. Y.Hao isolated cell nuclei and Y.Hao and A.A. prepared samples for snRNA-seq. E.D., Y.Y.C., and J.H. performed animal behavior tests. S.Y.Y. and A.Z. aided in cohort tissue collection. A.R., S.W., A.T, J.S, and I.L helped in developing and optimizing experimental procedures and conditions. T.O. and M.K. provided advice and guidance on the study. YH and MS supervised the study. YL, YH, and MS wrote the manuscript. All authors have read and approved the manuscript.

## Declaration of interests

Y.H. is a co-founder and scientific advisory board member of GABAeron Inc. All other authors declare no competing interests.

Patent on this work was filed in 2024. U.S. Application Serial No. 63/603,372

## Notes

### Competing Interest Statement

Dr Yadong Huang is a co-founder and scientific advisory board member of GABAeron Inc. All other authors declare no competing interests.

