## Supplemental Figures for "Data-driven discovery of cell-type-directed network-correcting combination therapy for Alzheimer’s disease"

Supplementary Table 1: Summary of cohort, sample, and feature sizes for individual and integrated datasets

| Study | Cohort size | Total cell number | Number of features |
|---|---|---|---|
| Mathys et al. | 24 AD, 24 ctl | 70,634 | 17926 |
| Zhou et al. | 11AD, 11 ctl | 72,489 | 33694 |
| Lau et al. | 12 AD, 9 ctl | 165,959 | 33538 |
| Integrated dataset (standardized and filtered case & ctl) | 37 AD, 29 ctl | 137,065 | 29120 |

Supplementary Table 2: Case-control standardization reference

| **CERAD** | | **Braak** | |
|---|---|---|---|
| score | Coding | score | Severity |
| 1 | Definite | V & VI | Dementia |
| 2 | Probable | III & IV | Mild symptoms |
| 3 | Possible | I & II | Asymptomatic |
| 4 | No AD | | |

Supplementary Table 3: AD vs control differentially expressed gene list (excel file)

Supplementary Table 4: AD outcome measures after drug exposure

| Patient data from UC-wide Electronic Medical Record | | | | | | | | |
|---|---|---|---|---|---|---|---|---|
| | relative risk scores | p value | total # of patients (exposed to drug) | # with AD diagnosis | AD % in drug group | AD % in propensity matched controls | drug indications | targeting cell types |
| Letrozole | 0.466 | 1.27E-37 | 10841 | 110 | 1.01 | 2.18 | breast cancer | Ex.Neu, In.Neu |
| Irinotecan | 0.195 | 9.83E-23 | 2227 | 8 | 0.004 | 0.019 | cancer | Ast, Mic, OPC |
| Methotrexate | 0.650 | 4.10E-12 | 27547 | 335 | 1.22 | 1.87 | cancer autoimmune | Ex.Neu, Ast |
| Ciclopirox | 0.927 | 0.0306 | 47642 | 1164 | 2.44 | 2.64 | skin infection, dermatitis | Ex.Neu, Ast, Opc |
| Sirolimus | 0.509 | 0.0209 | 4155 | 14 | 0.34 | 0.66 | immuno-suppression cancer | Ex.Neu, Ast, Oli |
| Etoposide | 0.885 | 0.711 | 1443 | 23 | 1.57 | 1.77 | cancer | Ex.Neu, Ast |
| Trifluoperazine | 0.842 | 0.835 | 197 | 8 | 0.039 | 0.046 | schizophrenia | Ex.Neu, Ast |
| Vorinostat | 0 | 0.178 | 59 | 0 | 0 | 4 | lymphoma cancer | Ex.Neu, Mic |
| Valproic acid | 5.818 | 0 | 30172 | 3619 | 11.99 | 2.06 | migraine seizure, bipolar | Ex.Neu, In.Neu, Ast |
| Haloperidol | 3.042 | 0 | 72636 | 5492 | 7.56 | 2.49 | schizophrenia | Ex.Neu, Ast |

Supplementary Table 5: Sex-stratified AD outcome measures after exposure of letrozole or irinotecan

| Patient data from UC-wide Electronic Medical Record | | | | | | | | |
|---|---|---|---|---|---|---|---|---|
| | sex | relative risk scores | p value | total # of patients (exposed to drug) | AD % in drug group | AD % in propensity matched controls | drug indications | targeting cell types |
| Letrozole | male | 0.444 | 0.170 | 325 | 0.012 | 0.027 | infertility | Neurons |
| Letrozole | female | 0.431 | 3.07E-19 | 10605 | 0.011 | 0.025 | breast cancer | Neurons |
| Irinotecan | male | 0.300 | 0.044 | 892 | 0.003 | 0.011 | cancer | Ast, Mic, OPC |
| Irinotecan | female | 0.154 | 0.044 | 623 | 0.002 | 0.010 | cancer | Ast, Mic, OPC |

Supplementary Table 6: Combination-treatment vs vehicle-treatment differentially expressed gene list (excel file)

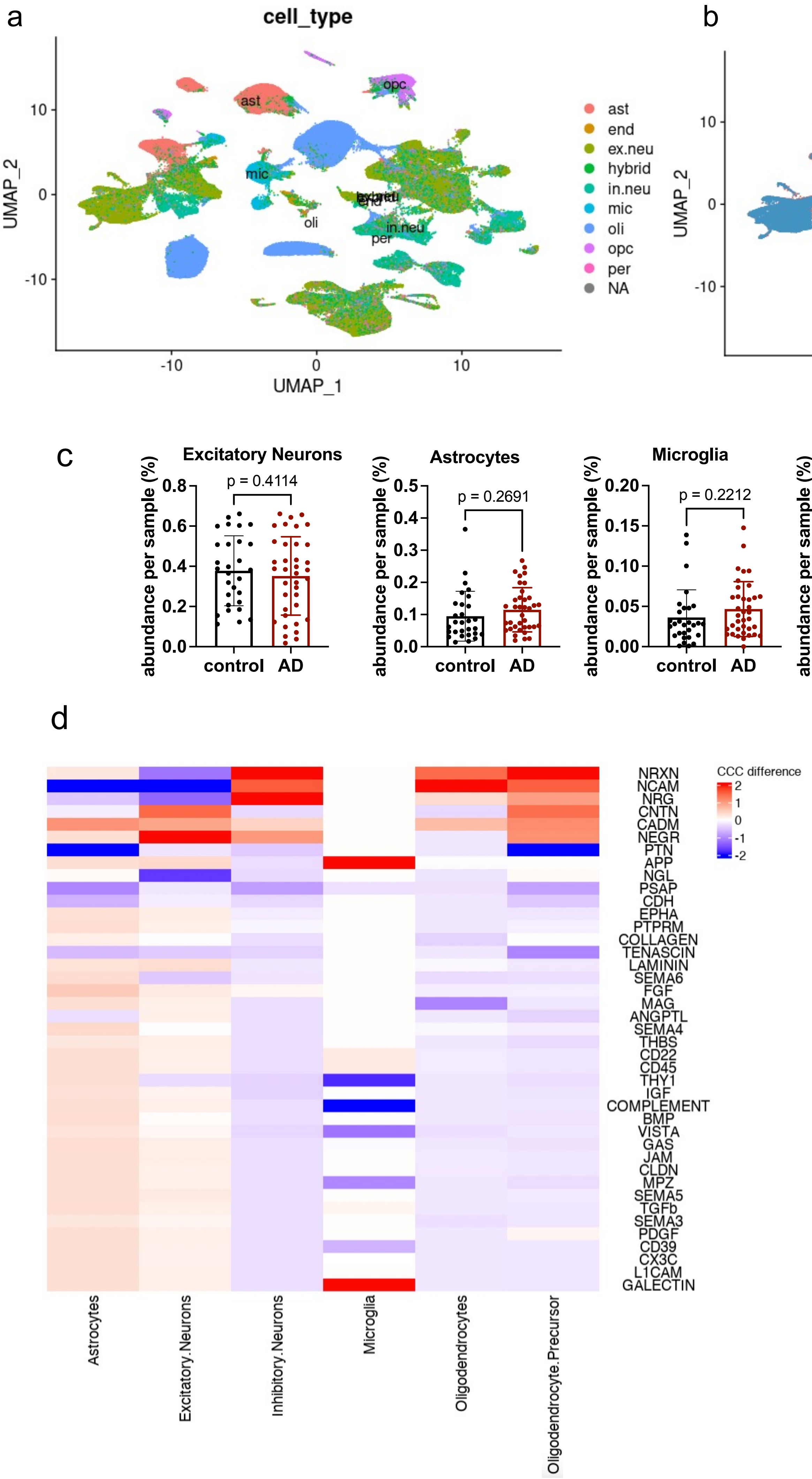


**Figure S 1a,b.** UMAP plot of merged samples before CCA integration, labeled with cell type identities (a) and disease status (b). **c.** Scattered plots of cell type abundance (percentage) for the other five cell types, comparing AD and controls (unpaired two-sided t-test). d. Heatmap displaying pathway-level cell-cell-communication (CCC) difference comparing AD and controls and stratified by cell type. Color value represents the scaled CCC probability difference in AD compared to controls. Signaling pathways represent ligand-receptor pairs based on the KEGG Pathway database.

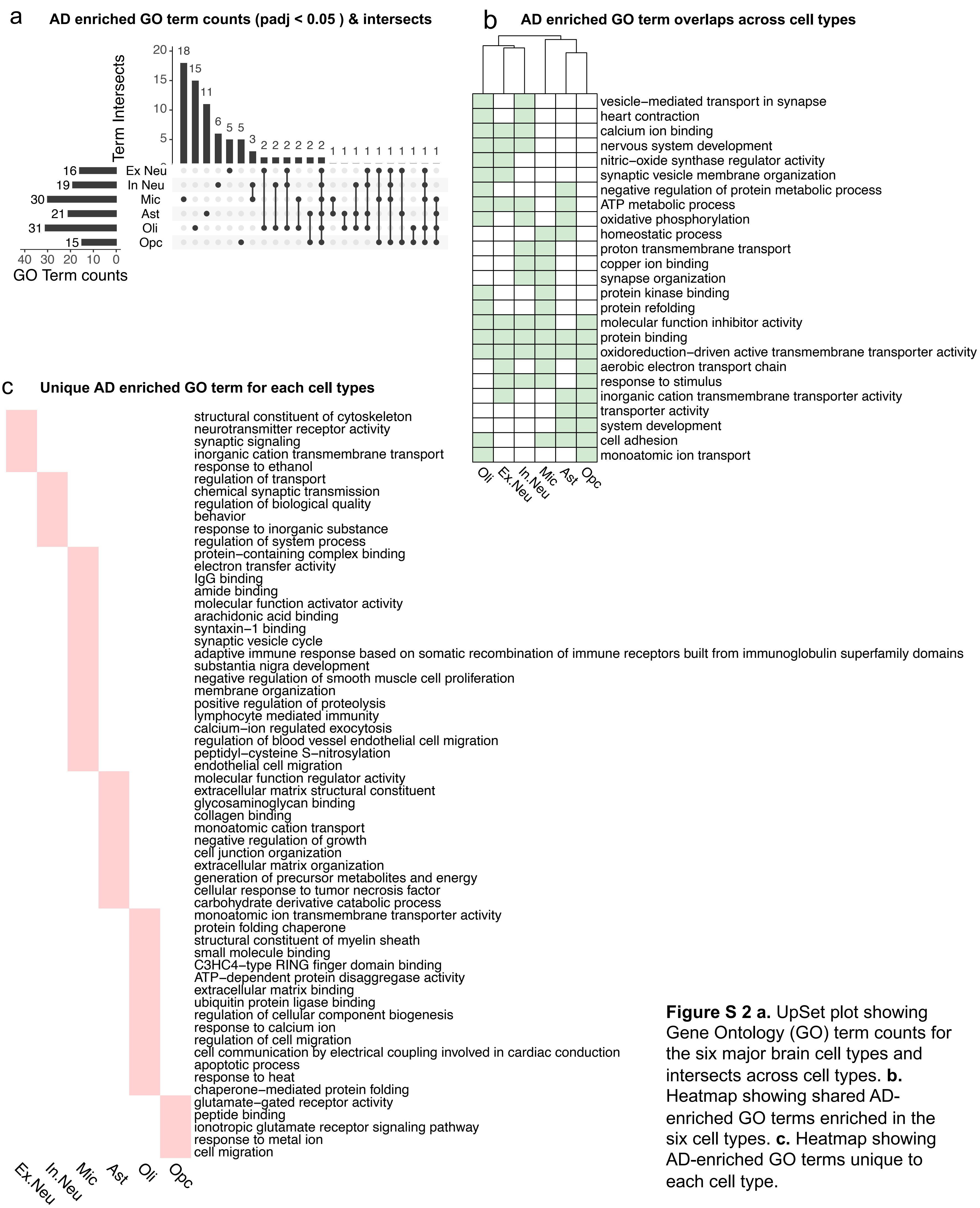


**Figure S 2 a.** UpSet plot showing Gene Ontology (GO) term counts for the six major brain cell types and intersects across cell types. **b.** Heatmap showing shared AD-enriched GO terms enriched in the six cell types. **c.** Heatmap showing AD-enriched GO terms unique to each cell type.

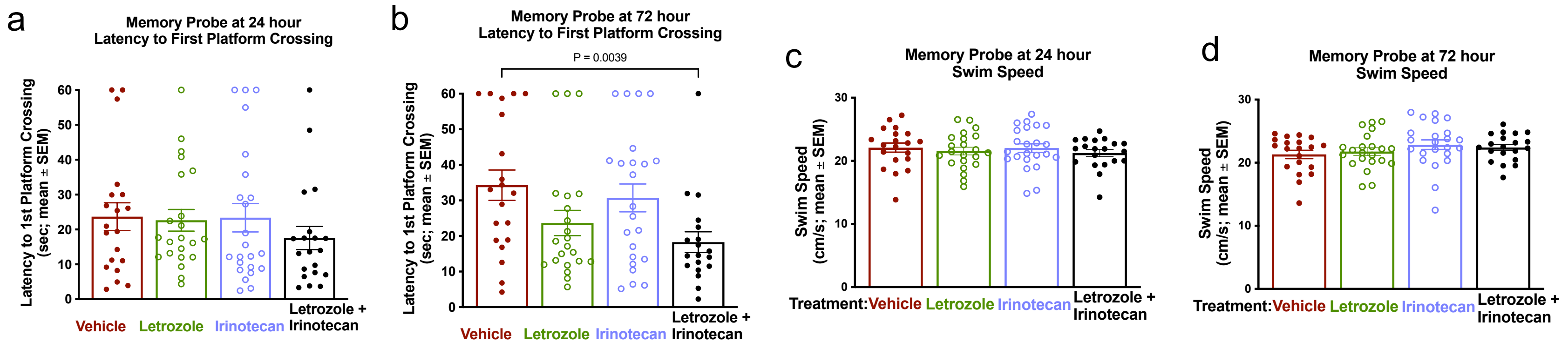


**Figure S 3. a,b.** Memory probes measuring latency to first time of platform crossing at 24 hour (a) and 72 hour (b) demonstrated long-term location memory rescue in combination treated mice only, at 72 hour after platform removed. **c,d.** Swim speed at 24hour (a) and 72 hour (b), no significant difference in swim speed across groups indicating that the observed behavioral differences were not confounded by visual or motor impairment. **a,b,c,d.** Ordinary one-way ANOVA were applied between all treatment groups and vehicle control group, p-values were adjusted by Dunnett's multiple comparison test.

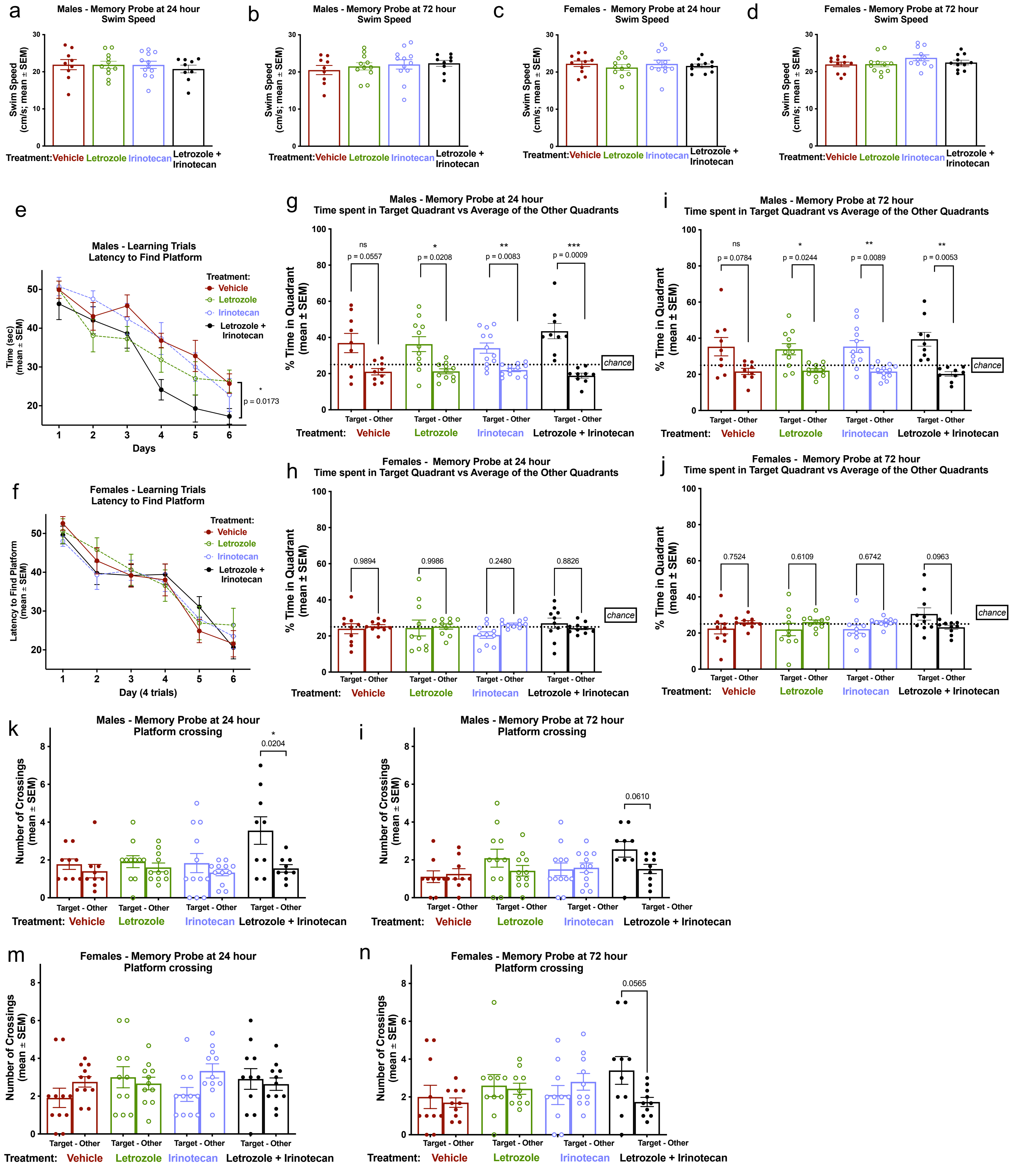


**Figure S 4. a,b,c,d.** Swim speed at 24hour (a,c) and 72 hour (b,d) in males (a,b) and in females (c,d), no significant difference in swim speed across groups indicating that the observed behavioral differences were not confounded by visual or motor impairments. **e,f**, Escape latency over learning days 1-6 in male mice (e) and in female mice (d) demonstrating significant better learning in combination-treated males. One-way repeated-measures ANOVA test was applied comparing vehicle group with treatment groups with Sidak's multiple comparison test. **g-j.** Memory probes of percent time spent in target quadrant vs average of other quadrant demonstrating a significant preference of the target quadrant by letrozole, irinotecan, and combination-treated males but not the vehicle-treated males at 24 hour (g) and 72 hour (i) after removing the hidden platform. No significant preference was observed in females at 24 hour (h) and 72 hour (j). Ordinary one-way ANOVA test was performed between target and other quadrants for each group, and p-value adjusted with Bonferroni multiple-comparisons testing. **k-n.** Memory probes measuring number of platform-location crossings in the target quadrant versus average of the other quadrants in males (k,l) and in females (m,n). Significantly more crossings in the target quadrant where the platform used to be only observed in the combination treatment group at 24 hour (k) in males, while there were a trend towards significant difference in males and in females at 72 hours (l,n) after the platform was removed. **a,b,c,d,k,i,m,n.** Ordinary one-way ANOVA were applied between all treatment groups and vehicle control group, p-values were adjusted by Dunnett's multiple comparison test.

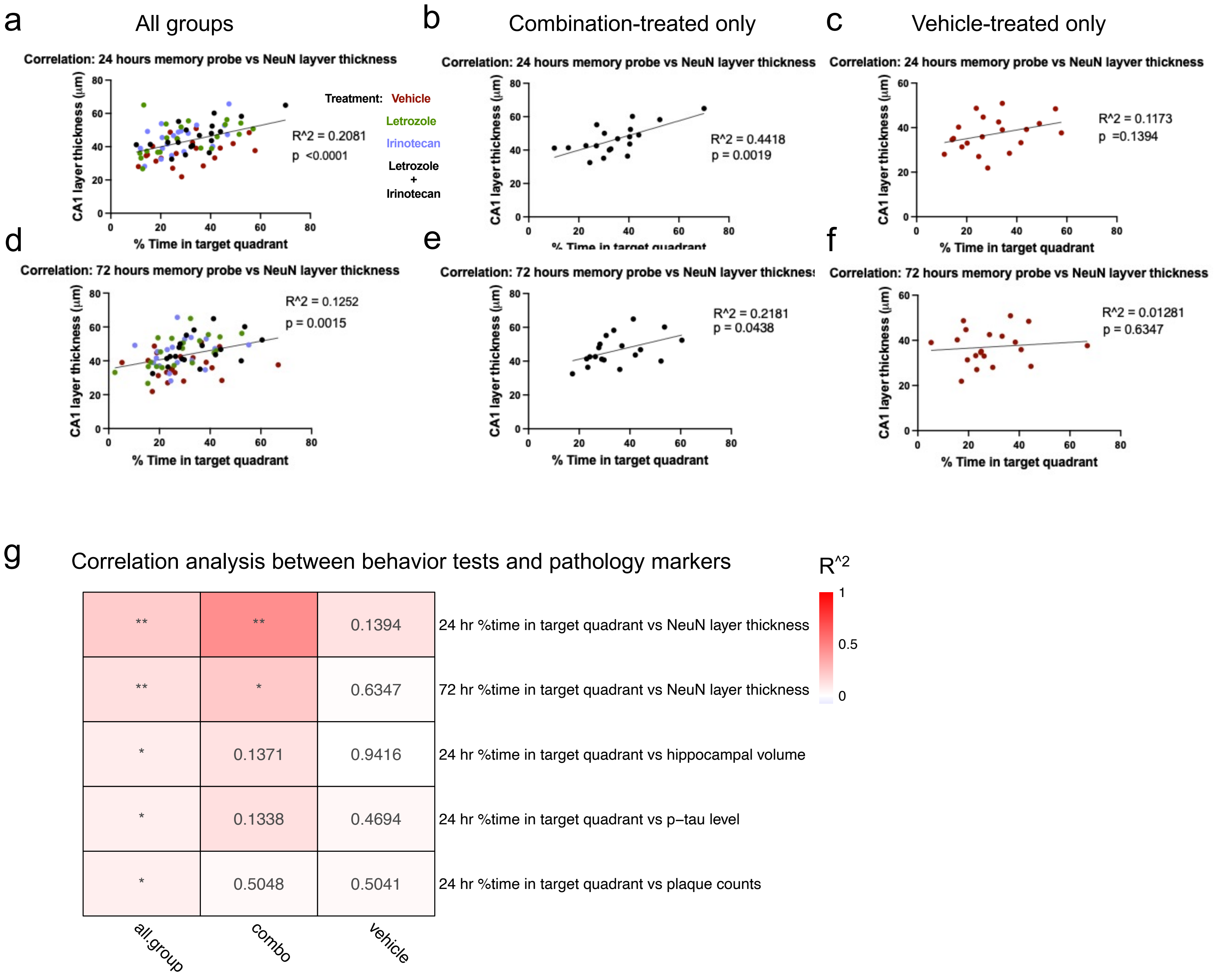


**Figure S 5 a-f,** Correlations between results from memory probes and pathological measures. Time spent in target quadrant at 24 hour versus NeuN layer thickness of entire experimental cohort (a) in combination treatment group only (b), in vehicle treatment group only (c). Time spent in target quadrant at 72 hour versus NeuN layer thickness of entire experimental cohort (d) in combination treatment group only (e), in vehicle treatment group only (f). g. Heatmap summarizing correlations between behavior test results and pathology measurements in all mice, in combination treated mice only, or in vehicle treated mice only. Color intensities resemble R-square values; number and * label p-values. * indicates p-value < 0.05, ** indicates p-value <0.01.

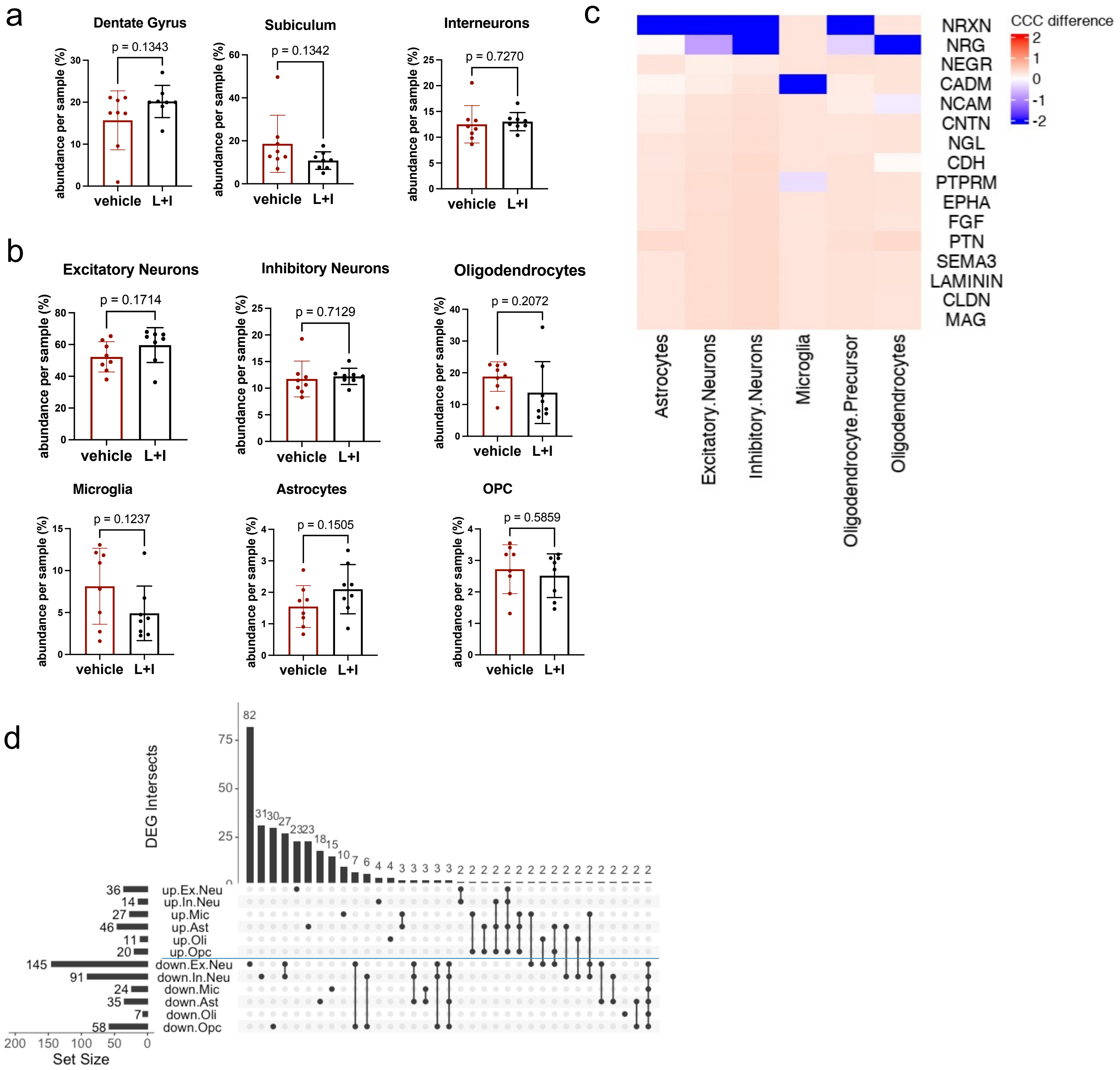


**Figure S 6 a.** Cell type abundance analysis for neuronal subtypes of hippocampus. **b.** Cell type abundance analysis for the six major brain cell types. **c.** Heatmap displaying pathway-level Cell-cell-communication difference between combination-treated and vehicle-treated and stratified by cell type. Value represents the scaled CCC probability difference in combination-treated compared to vehicle-treated. Signaling pathways represent ligand-receptor pairs based on the KEGG Pathway database. **d.** UpSet plot showing DEG counts (combination-treated compared to vehicle-treated, abs(LFC) > 0.1 & p-adj-value < 0.05) and intersect across groups (separated by cell types and direction of regulation).

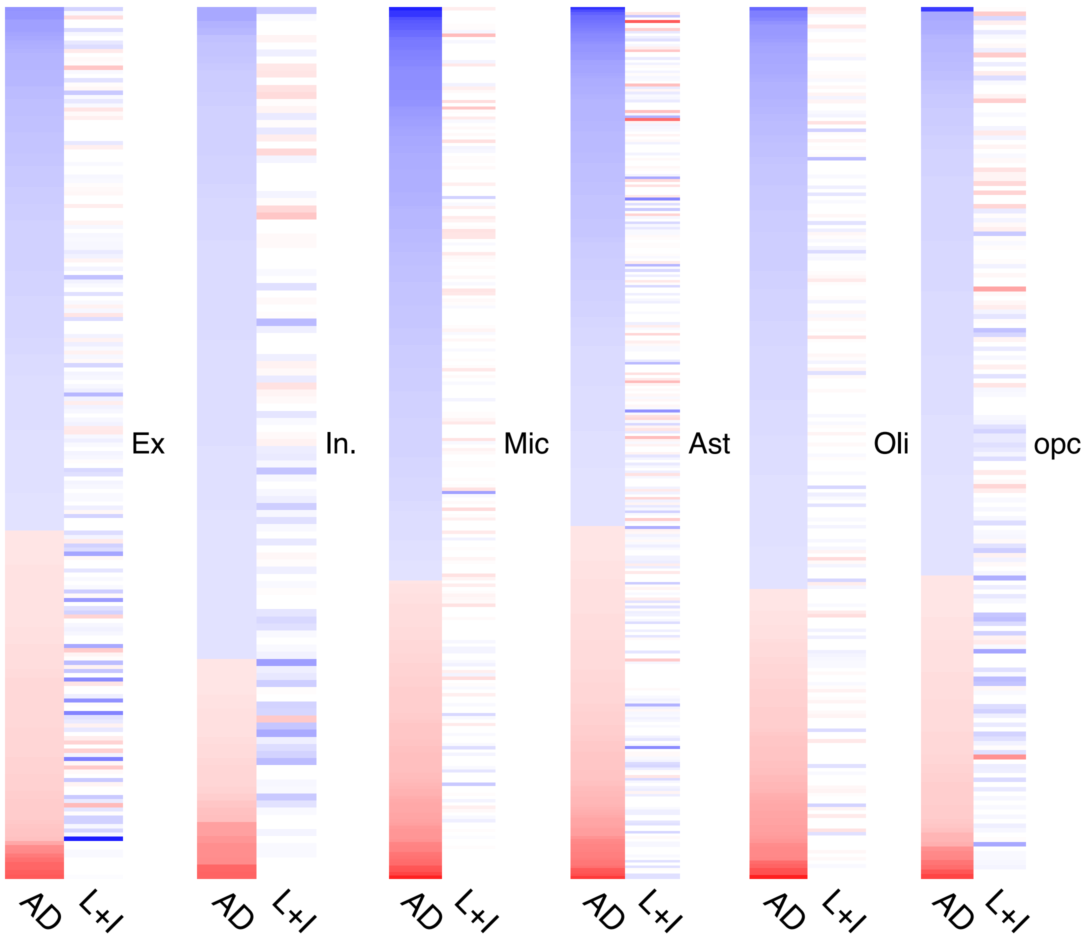


**Figure S7** All AD signature gene expression pattern comparison in human and after L+I treatment. Red indicates upregulation while blue indicates downregulation. White labels gene with no expression difference in treatment vs control or not expressed in mice.
